# β1 integrins regulate cellular behaviors and cardiomyocyte organization during ventricular wall formation

**DOI:** 10.1101/2023.08.28.555112

**Authors:** Lianjie Miao, Micah Castillo, Yangyang Lu, Yongqi Xiao, Yu Liu, Alan R Burns, Ashok Kumar, Preethi Gunaratne, C. Michael DiPersio, Mingfu Wu

**Author notes:** Correspondence to Mingfu Wu, Ph.D. Department of Pharmacological and Pharmaceutical Sciences, College of Pharmacy, University of Houston, Houston, TX 77204-5039.

## Abstract

**Aims:** The mechanisms regulating the cellular behavior and cardiomyocyte organization during ventricular wall morphogenesis are poorly understood. Cardiomyocytes are surrounded by extracellular matrix (ECM) and interact with ECM via integrins. This study aims to determine whether and how β1 integrins regulate cardiomyocyte behavior and organization during ventricular wall morphogenesis in the mouse.

**Methods and Results:** We applied mRNA deep sequencing and immunostaining to determine the expression repertoires of α/β integrins and their ligands in the embryonic heart. Integrin β1 subunit (β1) and some of its ECM ligands are asymmetrically distributed and enriched in the luminal side of cardiomyocytes, while fibronectin surrounds cardiomyocytes, creating a network for them. *Itgb1*, which encodes the β1 integrin subunit, was deleted via *Nkx2.5^Cre/+^* to generate myocardial-specific *Itgb1* knockout (B1KO) mice. B1KO hearts display an absence of trabecular zone but a thicker compact zone. The abundances of hyaluronic acid and versican are not significantly different. Instead, fibronectin, a ligand of β1, was absent in B1KO. We examined cellular behaviors and organization via various tools. B1KO cardiomyocytes display a random cellular orientation and fail to undergo perpendicular cell division, be organized properly, and establish the proper tissue architecture to form trabeculae. The reduction of Notch1 activation was not the cause of the abnormal cellular organization in B1KO hearts. Mosaic clonal lineage tracing shows that *Itgb1* regulates cardiomyocyte transmural migration and proliferation autonomously.

**Conclusions:** β1 is asymmetrically localized in the cardiomyocytes, and its ECM ligands are enriched in the luminal side of the myocardium and surrounding cardiomyocytes. β1 integrins are required for cardiomyocytes to attach to the ECM network. This engagement provides structural support for cardiomyocytes to maintain shape, undergo perpendicular division, and establish cellular organization. Deletion of *Itgb1*, leading to ablation of β1 integrins, causes the dissociation of cardiomyocytes from the ECM network and failure to establish tissue architecture to form trabeculae.

## Introduction

The heart is the first functional organ formed in mammalian embryonic development^1^. During cardiac morphogenesis, cardiac progenitor cells from the cardiac crescent migrate toward the ventral midline to form a linear heart tube with a smooth inner surface^2,3^. When the heart tube undergoes looping, the myocardium along the outer curvature of the tube grows inward and forms sheet-like structures that extend from the myocardium^2,3^, which are the newly initiated trabeculae^4,5^. As the early embryonic heart does not have a coronary circulatory system to perfuse itself, the sheet-like trabeculae function to increase surface area to facilitate nutrient and oxygen exchange. A lack of trabeculation will result in less availability of oxygen and nutrients in the myocardial tissue and end with embryonic lethality, while excess trabeculation will cause left ventricular non-compaction (LVNC) cardiomyopathy in humans^6–8^. Trabecular formation is a multistep process that includes, but is not limited to, trabecular initiation, specification, growth, and compaction^9–11^. Recent works show cardiomyocytes in the monolayer myocardium display polarity, and proper cellular behaviors such as oriented cell division and directional migration of the cardiomyocytes in the monolayer myocardium contribute to trabecular-initiation in an N-Cadherin dependent manner^11–15^. However, the underlying mechanisms regulating cell behavior and organization during trabecular morphogenesis and ventricular wall formation require further study. The cells in the monolayer myocardium are not typical cardiomyocytes as gap junction and intercalated discs are not fully formed yet. They present some features of epithelial cells as they display polarity and display squamous or cuboidal shape, but they are not epithelial cells, as these cells do not have the typical tight junctions and desmosomes^16^. So, how these cells become organized to establish the tissue architecture of trabeculae and ventricular wall is unknown.

Integrins are transmembrane heterodimeric proteins that consist of an α and a β subunit, each with a large extracellular domain, a single-pass transmembrane domain, and a cytoplasmic domain^17^. Integrins are the major cell surface receptors for adhesion to the ECM and mediate both inside-out and outside-in signal transduction pathways that control various cell functions, including proliferation, survival, migration, and gene expression ^17–19^. We found that β1 integrin subunit (β1), encoded by *Itgb1,* is highly expressed in the heart and is expressed in all cardiac cell types. Global deletion of *Itgb1* is lethal at the pre-implantation stage^20,21^. Endothelial-specific knockout of *Itgb1* disrupts cell polarity and arteriolar lumen formation^22^ and results in severe vascular defects and lethality at E10.5^23,24^. Cardiomyocyte-specific knockout of the *Itgb1* via *Mlc-2vCre* results in myocardial fibrosis and cardiac failure in the adult heart^25^, and *Itgb1* deletion via the transgenic *Nkx2.5Cre* line perturbs cardiomyocyte proliferation with progressive cardiac abnormalities seen toward birth^26^. Since both Cre lines ablate *Itgb1* at later stages, requirements for β1 integrins at earlier stages of trabecular and ventricular morphogenesis have been unknown. This study found that *Itgb1* deletion at an early stage via *Nkx2.5^Cre/+^* causes distinct but interesting defects.

The purpose of this study was to determine whether and how β1 integrins regulate cardiomyocyte behavior and organization during ventricular wall morphogenesis in the mouse. We conclude from our findings that cardiomyocyte-ECM interactions that are mediated by β1 integrins provide a structural framework and a microenvironmental niche to support the cellular orientation, cellular behaviors, cellular organization and ventricular specification, and that disruption of β1 integrins causes abnormal cell orientation, cellular behaviors and organization, and asymmetric cell division, resulting in trabeculation defects and ventricular wall specification defects. This study provides a basis for the pathogenesis of trabeculation defects and enhances our understanding of how ECM regulates ventricular wall morphogenesis.

## Results

### β1 integrins and their ligands are asymmetrically localized in the myocardium

Integrins are essential to mediate cell-ECM interactions^27^. A detailed expression repertoire of integrins and their ligands during cardiac morphogenesis is not available^27^. Therefore, we examined the expression of genes that encode α or β subunits of integrins and their ligands in hearts at E9.5 by mRNA deep sequencing, and their expression patterns by immunostaining and RNAScope (Table 1&Fig. 1A-F). Of all the genes that encode the β subunits, *Itgb1* is the most abundantly expressed (Table 1), and it is expressed in cardiomyocytes, endocardial cells, and epicardial cells based on immunostaining (Fig. 1A). Surprisingly, β1 is not evenly distributed on the membrane of cardiomyocytes but enriched in the luminal side of the membrane of most of the cardiomyocytes (Fig. 1A1&2, yellow arrows). It is also asymmetrically distributed in cardiomyocytes when the myocardial layer has two layers of cells (Suppl. Fig 1A. and Suppl. Movie 1). Of the genes that encode the α subunits, *Itga6* and *Itga5* are the two most abundantly expressed genes that encode α subunits (Table 1). Integrin α5 subunit (abbreviated as α5), encoded by *Itga5*, is expressed in both trabecular and compact cardiomyocytes (Suppl. Fig. 2A). In contrast, integrin α6 subunit (abbreviated as α6), encoded by *Itga6,* is enriched in trabecular cardiomyocytes at both mRNA and protein levels (Fig.1C, C1, C2&Suppl Fig.2C). Integrin ligands, including fibronectin (Fn), collagen, vitronectin, laminin, and vascular cell adhesion molecules, are expressed in the vasculature or epicardium^17^. In the embryonic heart at E9.5, Fn is abundantly expressed, *Col4a1* is the most abundant gene that encodes collagens, and *lama4*, *lamb1,* and *lamc1* are the most abundantly expressed genes that encode laminin α4, β1, and γ1 isoforms, respectively (Table 1). Fn, the ligand for α5β1, occurs in two main forms: plasma Fn, which can be detected by immunostaining with an anti-40K, and cellular Fn, which can be identified by the alternatively spliced domain A specific antibody. We found that the plasma Fn is abundantly expressed proximal to the basal cell layer and luminal side of the myocardium. Furthermore, Fn surrounds the individual cardiomyocytes, creating an ECM scaffold for cardiomyocytes (Fig. 1E). In contrast, the cellular isoform is expressed in the AV canal but not in the ventricles (Suppl. Fig. 2E). Laminin α4, β1 and γ1 (Laminin 411), a ligand for α6β1 integrin, is expressed in the basement membrane and asymmetrically enriched on the luminal side of the myocardium (Suppl. Fig. 1G), as is collagen IV (Suppl. Fig. 2I), which is highly expressed in the heart^28^.

**Figure 1:**
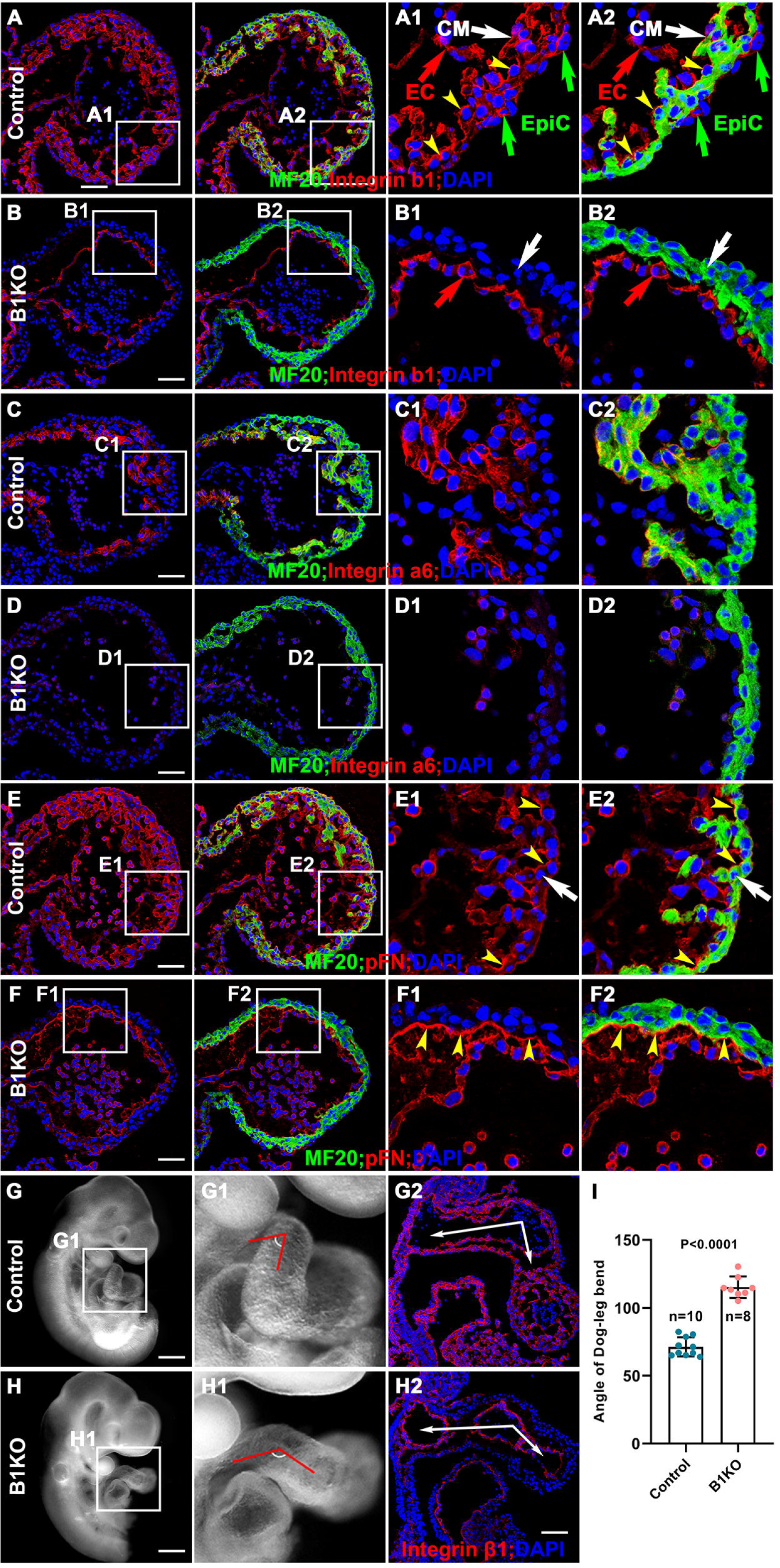
Expression of β1 integrin and its ligands in the E9.5 hearts. (A, A1&A2) In E9.5 hearts, β1 integrin is abundantly expressed in cardiomyocytes (CM), endocardial cells (EC), and epicardial cells (EpiC) and displays an asymmetrical distribution pattern being enriched in the luminal side of the cardiomyocytes. (B, B1&B2) *Itgb1* was efficiently deleted from the myocardium by *Nkx2.5^cre/+^*, confirmed by β1 integrin immunostaining. (C, C1&C2) Integrin α6 is enriched in trabecular cardiomyocytes. (D, D1&D2) Integrin α6 is reduced or absent in B1KO hearts. (E, E1&E2) Plasma Fn is abundantly expressed in the basal membrane and luminal side of the myocardium. (F, F1&F2) Plasma Fn is reduced in the myocardium of B1KO hearts, but remains expression in luminal side of the myocardium. (G-I) The B1KO hearts display an abnormal OFT morphogenic defect, with a larger rotation angle at the Dog-Leg bend. Scale bar: 50μm in A-F, 300μm in G&H.

**Figure 2:**
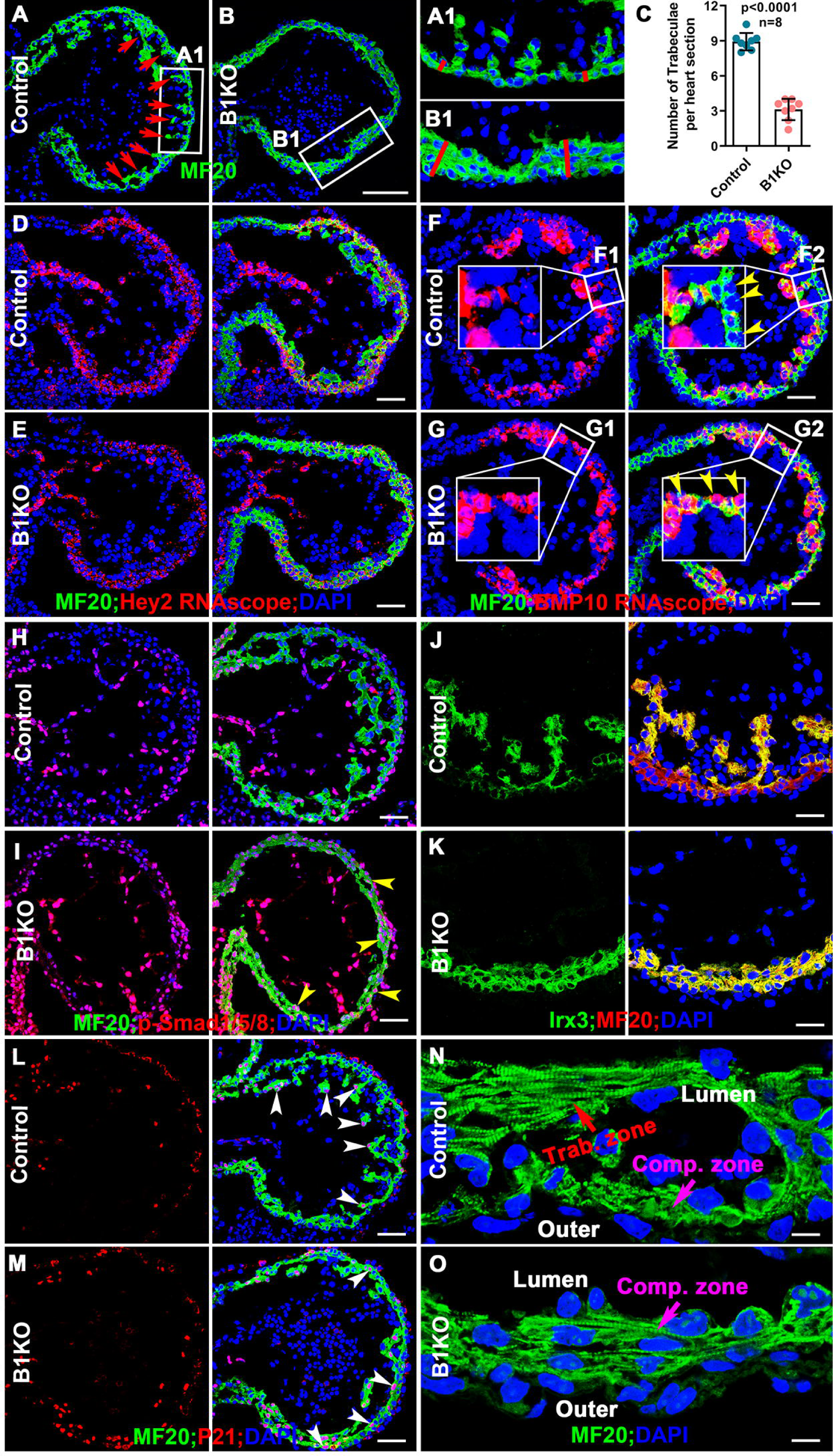
B1KO hearts display defects in trabecular formation and trabecular/compact zone specification. (A-C) The B1KO hearts display a trabeculation defect, with a significantly smaller number of trabeculae per section and a thicker ventricular wall. (D&E) *Hey2*, a specification marker for the compact zone, is highly enriched in the compact zone of both control and B1KO hearts. (F&G) In the control hearts, *BMP10*, a specification marker for the trabecular zone, is highly enriched in the trabecular cardiomyocytes but not compact cardiomyocytes. In the B1KO hearts, *BMP10* is also highly enriched in some compact cardiomyocytes, in addition to the trabecular cardiomyocytes. (H&I) Smad1/5/8 showed more phosphorylation in cardiomyocytes of the B1KO hearts, indicated by p-Smad1/5/8, a readout for BMP10. (J&K) Irx3, a trabecular cardiomyocyte marker, is enriched in the trabecular zone in control hearts but is expressed in the compact zone in B1KO hearts. (L&M) P21 is mainly expressed in the trabecular cardiomyocytes but is also highly expressed in the compact cardiomyocytes in the B1KO hearts. (N&O) The trabecular cardiomyocytes, sarcomeric array formation, are more organized than the compact cardiomyocytes in control hearts. In the B1KO hearts, the compact cardiomyocytes formed a sarcomeric array, indicating an early differentiation defect. Scale bar: 50μm in A-M, 5μm in N&O.

**Table 1.** Expressions of Integrins and their ligands in E9.5 hearts.

| symbol | baseMean | log2FoldChange | pvalue |
| --- | --- | --- | --- |
| Itga6 | 15496.92185 | 0.335664289 | 4.18E-10 |
| Itga5 | 4807.254709 | -0.061236291 | 0.20244954 |
| Itgav | 2142.962146 | -0.038447686 | 0.707961797 |
| Itga4 | 1876.231852 | -0.252868185 | 0.000496303 |
| Itga9 | 1824.099191 | -0.046879344 | 0.53081939 |
| Itga1 | 1275.069268 | -0.245572751 | 0.002105139 |
| Itga3 | 737.6016552 | -0.150938337 | 0.138973659 |
| Itga2b | 430.4990634 | 0.126087859 | 0.495450734 |
| Itga10 | 155.2980159 | 0.160179897 | 0.381722821 |
| Itga8 | 141.1315812 | -0.47723196 | 0.050037637 |
| Itga2 | 61.24040399 | 0.021476907 | 0.942211481 |
| Itga7 | 48.70622252 | -0.832033757 | 0.046689928 |
| Itgae | 36.1021102 | -0.017610151 | 0.961376462 |
| Itgal | 20.64407217 | 0.272856541 | 0.589842947 |
| Itga11 | 20.61786085 | -0.094726064 | 0.852970763 |
| Itgad | 14.23869144 | 0.102638633 | 0.863373361 |
| Itgam | 9.445725768 | 0.550455168 | 0.449605271 |
| Itgax | 1.067665895 | -2.046504706 | 0.404620739 |
| Itgb1 | 13097.54951 | -0.806979211 | 3.20E-84 |
| Itgb1bp2 | 1680.649481 | 0.059181393 | 0.390070033 |
| Itgb3 | 1244.226623 | -0.215286178 | 0.008090809 |
| Itgb5 | 1234.860529 | 0.097639746 | 0.227267051 |
| Itgb1bp1 | 343.9195277 | -0.090082488 | 0.50564828 |
| Itgb3bp | 341.835156 | -0.099062994 | 0.442618139 |
| Itgb8 | 264.9667078 | -0.093060695 | 0.522954196 |
| Itgb2 | 38.12422731 | 0.367946672 | 0.35432961 |
| Itgb4 | 26.39955801 | 0.228166267 | 0.66007866 |
| Itgb7 | 4.529116186 | 1.173006406 | 0.289827753 |
| Itgb6 | 2.100802743 | 0.346116136 | 0.825130566 |
| Itgb2l | 0.921997253 | 0.062760271 | 0.981310739 |
| Lamc1 | 11137.66658 | -0.008979321 | 0.836020299 |
| Lamb1 | 8290.679242 | 0.058119707 | 0.219516928 |
| Lama4 | 5624.804458 | -0.002547953 | 0.955238015 |
| Lama5 | 3268.083023 | -0.17772571 | 0.016466224 |
| Lamb2 | 3101.234381 | 0.200027277 | 0.036551497 |
| Lama1 | 452.468331 | -0.014017722 | 0.89969672 |
| Lama2 | 301.5820561 | 0.05231657 | 0.697809472 |
| Lamc2 | 145.9615107 | -0.554639512 | 0.005402435 |
| Lamb3 | 78.32032565 | 0.260448715 | 0.425521045 |
| Lama3 | 41.49436877 | -0.165902076 | 0.640995185 |
| Lamc3 | 19.21278617 | -0.937737738 | 0.122270688 |
| Fn1 | 54378.72786 | 0.036682721 | 0.312994843 |
| Vcam1 | 3191.127312 | 0.185906499 | 0.002174042 |
| Vtn | 6.65353464 | -3.457998104 | 0.00614626 |
| Col4a1 | 20866.09636 | -0.031527466 | 0.465346995 |
| Col2a1 | 14377.64281 | -0.469676661 | 3.62E-10 |
| Col18a1 | 13285.87809 | -0.09264386 | 0.107736858 |
| Col4a2 | 12524.17741 | -0.031760744 | 0.531309141 |
| Col3a1 | 5697.927173 | -0.391217463 | 2.89E-06 |
| Col4a5 | 5229.892031 | -0.075922729 | 0.194680351 |
| Col5a1 | 5048.677797 | 0.055519868 | 0.435512861 |
| Col5a2 | 3061.962381 | -0.103096736 | 0.055056056 |
| Col1a2 | 1992.371686 | 0.061368255 | 0.363804442 |
| Col4a6 | 1646.848477 | -0.17155072 | 0.013045162 |

### β1 integrins are required for the expression of Fn and α6

To study the functions of β1 integrins in cardiac morphogenesis, *Itgb1*, encoding β1 was deleted via the *Nkx2.5^Cre/+^*to generate *Nkx2.5^Cre/+^*; *Itgb1^fl/fl^* (B1KO). The deletion of *Itgb1* in the myocardium in B1KO was confirmed by the absence of β1 in the myocardium but not in endocardial cells (Fig. 1B). β1 heterodimerizes with α5, encoded by *Itga5*, or α6, encoded by *Itga6*, to bind to Fn and laminin 411, respectively^17^. As already mentioned, the genes that encode β1, α5 or α6 integrin subunits, or their ligands are abundantly expressed in the heart at E9.5 (Table 1). We therefore examined if *Itgb1* deletion affects the expression of α5 or α6 subunits and/or their ligands via mRNA deep sequencing and immunostaining. The transcript levels of genes that encode α and β integrins and ligands were not significantly different between control and B1KO hearts at E9.5 for any genes whose expression BaseMean is larger than 50 based on mRNA deep sequencing (Table 1). Immunofluorescence revealed that while localizations of α6 and Fn to the myocardium (Fig. 1C-F) were both significantly reduced or absent in B1KO hearts, Fn deposition to the luminal side of the myocardium was not affected, suggesting that endocardial cells can deposit Fn to the cardiac jelly (Fig. 1E). The Fn surrounding individual cardiomyocytes creates an ECM scaffold in the myocardium in control, while this ECM scaffold was absent in B1KO. These data indicate that β1 is required for the stabilization of both α6 and Fn in the myocardium. Other integrin subunits, such as α5, and other ECM ligands, such as collagen IV and laminin, did not display an obviously different pattern in B1KO compared with the controls (Suppl. Fig. 2A&B, G-J). The B1KO embryos died before E11.5 (Table 2), a more severe phenotype than the published transgenic Cre line *Nkx2.5Cre* mediated *Itgb1* knockouts^26^, in which *Itgb1^fl/fl^*was deleted at a later developmental stage, and the *Mlc-2vCre* mediated knockout, which results in myocardial fibrosis and cardiac failure in the adult heart^25^. The B1KO hearts display an abnormal OFT morphogenesis with a significantly large angle of Dog-Leg (Fig. 1G-I). The expression of cardiomyocyte marker MF20 shows slightly different patterns from the control hearts at E9.5, as the cardiac progenitor cells close to the endoderm express MF20, a marker for cardiomyocytes, in B1KO but not in the control heart (Suppl. Fig. 3A&B). These data suggest a defect in cardiac progenitor cell differentiation, consistent with a previous report of *Itgb1* in cardiac progenitor cell differentiation^29^ and a study showing *Itgb1* null embryonic stem cells show an increased differentiation of cardiac cells in the *in vitro* system^30^. In summary, considering that Fn is required for heart morphogenesis^31^, the absence of α6 and Fn in B1KO hearts suggests that β1 integrins mediate multiple and distinct integrin-ECM interactions that are collectively essential for heart morphogenesis.

**Table 2.** Survival rate.

| **Age** | **Total** | **KO** | **Harvested/Expected percentage of KO** |
| --- | --- | --- | --- |
| E8.5 | 165 | 41 | 25/25 |
| E9.5 | 483 | 105 | 22/25 |
| E10.5 | 144 | 28 | 19/25 |
| E11.5 | 25 | 1 | 4/25 |
| E12.5 | 12 | 0 | 0/25 |

### B1KO hearts display defects in trabecular formation and trabecular/compact zone specification

Trabeculae are sheet-like structures extending from the myocardium to the heart lumen and function to increase surface area when the coronary system is not yet established^32^. The B1KO hearts display a significantly smaller number of trabeculae per section examined at E9.5, suggesting a trabeculation defect (Fig. 2A-C); however, the compact zone of B1KO is 2.4 times thicker than the control (Fig. 2A1&B1). The thicker compact zone but reduced number of trabeculae suggests that cardiomyocytes failed to be properly organized and redistributed between trabecular and compact zones. Trabecular cardiomyocytes, which take the major responsibility for pumping at the early stages of cardiac development, are more differentiated than compact zone cardiomyocytes^4^. Trabecular and compact cardiomyocytes display different features, with trabecular cardiomyocytes exhibiting a lower proliferation rate and being more molecularly mature than cardiomyocytes of the compact zone^33^. For example, *p21*, *Irx3, BMP10,* Sphingosine 1-phosphate receptor-1, and *Cx40* are highly expressed in the trabecular zone, while *Tbx20*, *Hey2,* and *N-Myc* are highly expressed in the compact zone^4,9,12,15,34–36^. The regulation of their specific expression patterns in trabecular and compact zones has not been elucidated. We wished to determine if β1 regulates trabecular/compact zone specification by examining the expression patterns of trabecular and compact zone markers in B1KO hearts. *Hey2* is expressed in the compact zone in both control and B1KO (Fig. 2D&E), but the abundance of *Hey2* mRNA in the compact zone in B1KO was reduced compared to control littermates (Fig. 2D, E& Suppl. Fig. 3C). *Bmp10*, a marker for trabecular cardiomyocytes, is expressed in the trabecular cells in the control (Fig. 2F, F1&F2), but is highly expressed in compact cells in B1KO (Fig. 2G, G1&G2). B1KO hearts display increased staining for *Bmp10* compared to the control (Fig. 2F, G) and mRNA expression is increased (Suppl. Fig. 3C). The increased staining for *Bmp10* in B1KO is supported by the increased staining for Smad1/5/8 phosphorylation in cardiomyocytes (Fig. 2H&I), a readout for *BMP10*^37^. *Irx3* is expressed in the trabecular zone in the control as anticipated but is expressed in the compact zone in B1KO (Fig. 2J&K). *p21* was mainly expressed in the trabecular cardiomyocytes in control but staining in compact cardiomyocytes was greater in B1KO (Fig. 2L, M &Suppl. Fig. 3D). The abnormal expression patterns of trabecular and compact zone-specific markers in B1KO hearts suggest a trabecular and compact zone specification defect and implicate β1 integrins in the regulation of ventricular wall specification.

Since the trabecular cardiomyocytes are more mature than cardiomyocytes of the compact zone^33^, the specification defects in B1KO suggested the B1KO hearts may show an early maturation defect. We examined sarcomere organization in control and B1KO hearts at E9.5. The sarcomeric array formation in the trabecular zone of a control heart is more organized than that of the compact zone in a control heart (Fig. 2N). The sarcomeric array formation in the compact zone in B1KO is more organized than that of the control, but less organized than cells in the trabecular zone of the control (Fig. 2N&O). This was further confirmed by electron microscopy (EM), as only the trabecular zone displayed regular sarcomeric array in control, while the compact zone showed sarcomeric array in B1KO (Suppl. Fig. 3E&F), suggesting an early maturation defect.

### Genes involved in cell-ECM interactions, but not genes involved in adherens junction-mediated cell-cell interactions, are altered in B1KO hearts

Previous studies show that N-Cadherin, the major component of adherens junctions in the embryonic heart, regulates cellular behaviors and is required for trabecular initiation and ventricular wall morphogenesis^38,39^. We, therefore, examined the adherens junctions among cardiomyocytes and found that N-Cadherin localization was not affected by *Itgb1* deletion based on immunostaining (Fig. 3A&B), and the total N-Cadherin protein level was not significantly different between the control and B1KO samples (Fig. 3C). Ultrastructural observations suggest cell-cell contacts were not obviously affected (Fig. 3D&E).

**Figure 3:**
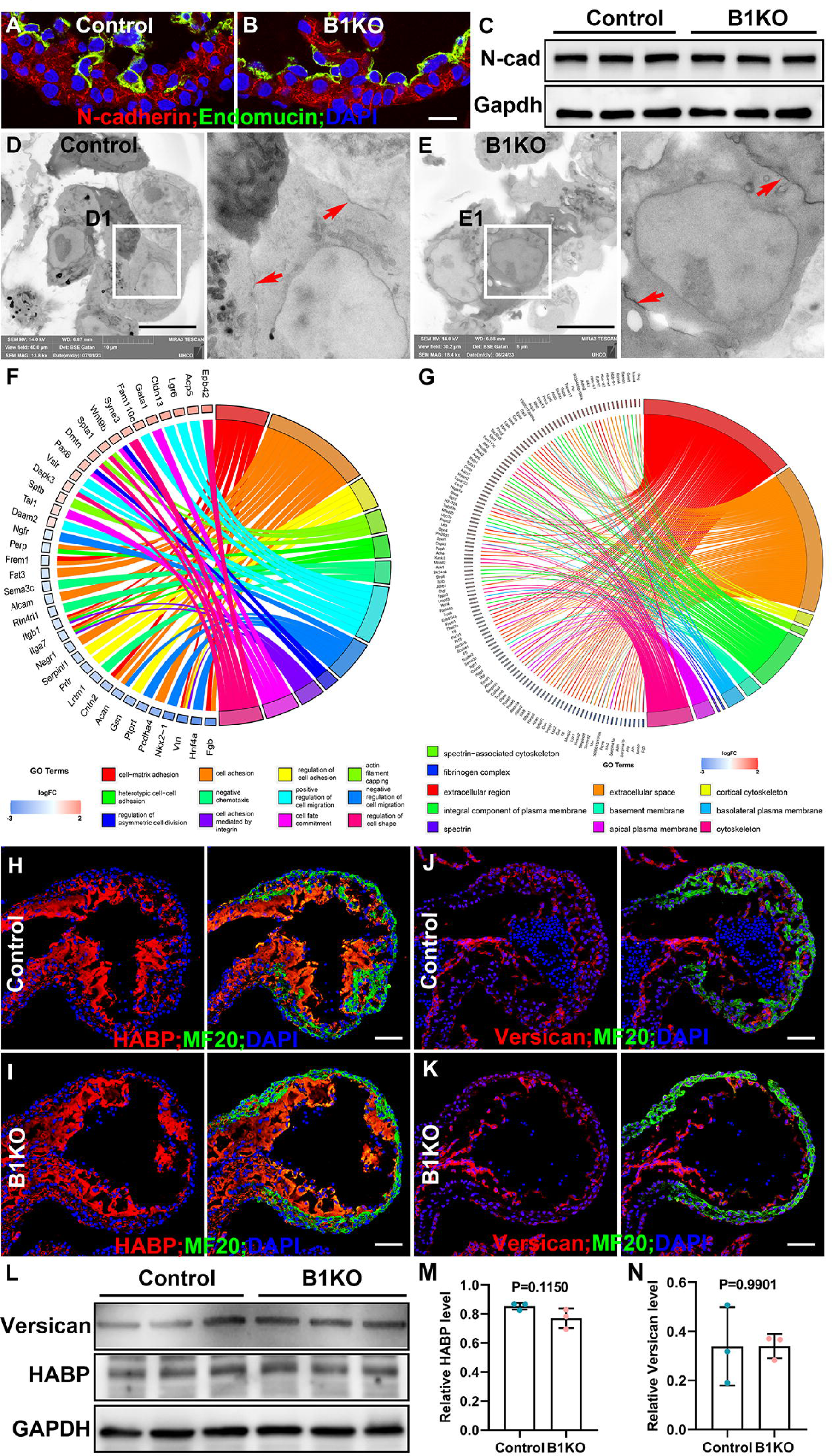
Cell-ECM interactions but not adherens junction mediated cell-cell interactions were affected. (A-C) Adherens junction was not changed in the E9.5 B1KO hearts compared with control hearts, indicated by N-cadherin expression pattern and protein level. (D&E) Based on electron microscopy, cell-cell junctions indicated by red arrows in D1 and E1 didn’t show an obvious difference between control and B1KO hearts. (F&G) Go analysis from mRNA deep sequencing showed the affected cellular components and biological processes in B1KO hearts. (H-N) The major cardiac jelly components, HABP and versican, didn’t show a significant difference between control and B1KO hearts based on immunostaining and Western Blot. Scale bar: 10μm in A&B, D&E, 50μm in H-K.

We next examined the transcript levels of genes involved in cell-ECM interaction, cell-cell interaction, and cardiac jelly via mRNA deep sequencing. Three E9.5 control or B1KO hearts were combined as one sample, and 3 samples of both control and B1KO were subjected to mRNA deep sequencing. Compared with control hearts, B1KO hearts showed significant differences in genes that encode cellular components of the extracellular region, extracellular space, basement membrane, fibrinogen complex, spectrin, and other cellular components (Fig. 3F). Further biological process analyses show that cell-matrix adhesion, heterotypic cell-cell adhesion, regulation of asymmetric cell division, cell migration, cell shape, etc., were significantly different between control and B1KO hearts (Fig. 3G). The expression of several matrix genes (*Has1*, *Has2*, *Has3*, and *Vcan*) was not altered in B1KO hearts based on mRNA deep sequencing. Major components in cardiac jelly, including versican and hyaluronic acid binding protein (HABP), were also not significantly different examined by immunostaining and western blot (Fig. 3H-N). Together, these findings indicate that the trabeculation defect in B1KO is not due to changes in genes that contribute to the cardiac jelly or cell-cell adherens junctions, but is more likely due to cell-ECM mediated interactions, which create a supportive niche for cardiomyocytes and might regulate ventricular specification.

### Cardiomyocytes in B1KO hearts display abnormal cellular organization

β1 is required for cardiomyocyte survival in a mosaic model^40^. To determine whether cell death causes the trabeculation defects in B1KO, control and B1KO hearts sections were stained for an apoptotic marker, cleaved Caspase 3. We found that the B1KO hearts displayed more cleaved caspase 3 positive cells in the OFT region, but not in the ventricular region of either control or B1KO hearts (Fig. 4A&B), suggesting that cell death in the ventricular region is not the cause of trabeculation defects. The proliferation rate was evaluated via BrdU pulse-labeling for one hour. The cardiomyocytes in B1KO hearts display a significantly reduced proliferation rate in both trabecular and compact cardiomyocytes (Fig. 4C-F). However, the reduction of the trabecular zone accompanied by a thickening of the compact zone (Fig. 2A) in B1KO hearts suggests that failed trabecular formation is not caused by reduced cell proliferation. Instead, the reduction or absence of trabeculae in B1KO is most likely caused by abnormal cellular organization and distribution between trabecular and compact zones.

**Figure 4:**
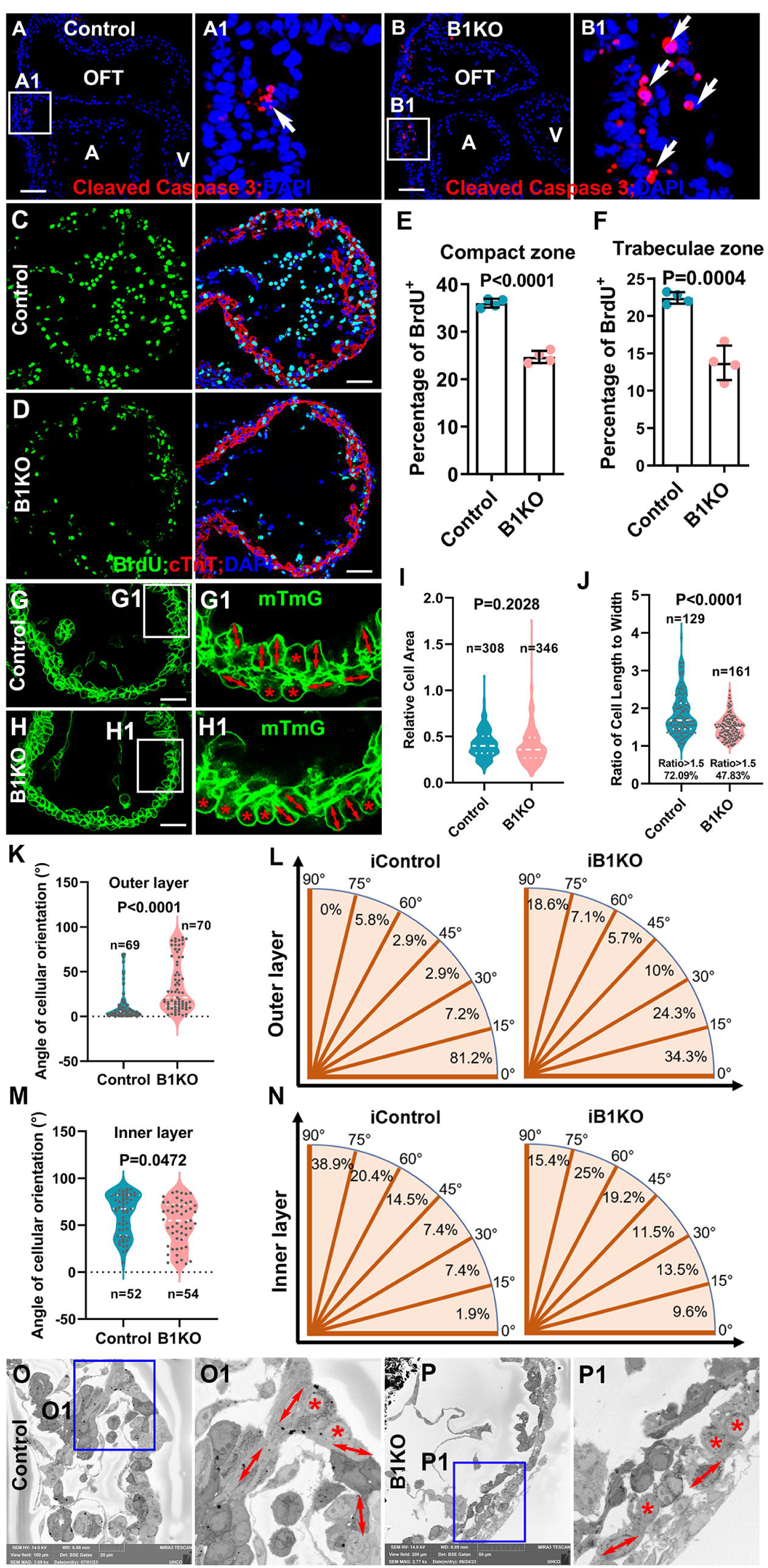
Cardiomyocytes in B1KO display abnormal cellular organization. (A&B) Immunostaining in the E9.5 B1KO hearts for cleaved caspase 3 reveals that B1KO hearts have more cell death in the OFT but not in the ventricular region compared with control hearts. (C-F) Based on BrdU pulse labeling, the cardiomyocytes in E9.5 B1KO hearts displayed lower proliferation rates than the control. (G&H) The mTmG reporter outlined cardiomyocytes in control and B1KO hearts at E8.75, and their cell size and shape were measured. (I) Cardiomyocyte size between control and B1KO hearts was not an obvious difference. (J) The lower percentage of cardiomyocytes display an orientation in B1KO hearts, with a ratio of cell length to width smaller than 1.5. (K-N) Oriented angles of cardiomyocytes in both control and B1KO hearts were measured (K&M). In the control hearts, most of the cardiomyocytes in the outer layer of the compact zone are oriented parallel to the heart wall, and cardiomyocytes in the inner layer of the compact zone are oriented perpendicularly to the heart wall. In the B1KO hearts, most of the cells display no or parallel orientation to the heart wall (L&M). (O-P) The abnormal cellular orientation in B1KO was confirmed by the EM. Scale bar: 50μm in A-D, 20μm in G&H.

The cells in the myocardium at early embryonic stage are not mature cardiomyocytes. How these cells become organized to form trabeculae is unknown. We hypothesize that the ECM scaffold provides a structural framework for cardiomyocytes to be properly organized and supports cellular shape, oriented cell division and migration. We, therefore, measured and compared the cell size by measuring cellular areas of cardiomyocytes between control and B1KO hearts. Specifically, the cell areas of individual cells in multiple sections were measured based on the membrane identified by membrane-localized GFP or RFP. We found that cell size is not significantly different between the control and B1KO at about E8.75 when the myocardium contains two layers of cells (Fig. 4G-I). We next examined the cell shape and cellular orientation of cardiomyocytes in the inner layer and outer layers of the myocardium. The ratio of length to width of cardiomyocytes was measured and quantified in both control and B1KO hearts. A cell with a ratio larger than 1.5 is defined as an oriented cell, while those less than 1.5 were quantified as non-oriented cells. The cellular orientation was measured by the angle between the heart surface and the axis of the oriented cells, as described previously^12^. We found that more than 72% of the control cells with a ratio larger than 1.5, while less than 48% of the cells in B1KO display a ratio larger than 1.5 (Fig. 4J).

A previous study showed that most myocardial cells in the outer layer display a parallel pattern, while cells in the inner layer display a perpendicular pattern^38^. We found that most cells (81.2%) in the outer layer display a parallel orientation with an angle of less than 15 (Fig. 4K&L). In contrast, the cells in B1KO heart display a random orientation with a significantly larger angle (Fig. 4K&L). We also measured the orientation of cells in the inner layer and found that 59.3% of the cells oriented with an angle larger than 60 degrees, and 9.3% of the cells oriented with an angle less than 30 degrees in control hearts (Fig. 4M&N), suggesting a perpendicular orientation pattern. In contrast, in B1KO hearts 40.4% of the cells are oriented with an angle larger than 60 degrees, and 23.1% of the cells are oriented with an angle less than 30 degree (Fig. 4M&N), suggesting a random orientation pattern in B1KO hearts. The abnormal cellular orientation in B1KO hearts was further confirmed by EM (Fig. 4O, O1, P, P1). The random orientation of the cells suggests a disrupted cellular organization and a loss of growth direction toward the heart lumen.

### β1 integrins are required for cardiomyocyte-oriented cell division and asymmetric cell division

Previous studies demonstrated that both oriented cell division (OCD) and directional migration contribute to trabecular initiation and trabecular specification in mice, which differs from the mechanism of apical constriction-mediated directional migration contributing to trabecular initiation in zebrafish ^11,12,16,41,42^. We asked whether β1 integrins are required for trabecular initiation. We first examined if β1 was asymmetrically distributed in dividing cardiomyocytes by staining for β1 and acetylated α-Tubulin to identify the mitotic spindles. We found that β1 is enriched to the membrane to the luminal side of the perpendicular dividing cells at all cell cycle stages (Fig. 5A&Suppl. Fig.1A) and is localized to the luminal side of both daughter cells in parallel dividing cells (Fig. 5B). This asymmetric distribution of β1 suggests that β1 integrins might be required for OCD and asymmetric cell division. To determine if the cardiomyocytes in B1KO display defective OCD, ∼E8.75 control and B1KO hearts were stained for acetylated α-Tubulin and P120 (an adherens junction-associated protein that marks the cardiomyocyte membrane). We measured the spindle orientations of mitotic cardiomyocytes of the left ventricular myocardium, and only the cells at anaphase or early telophase with both centrosomes in the same focal plane were quantified^12,43^. We found most mitotic cells displayed either perpendicular orientation (44%, n = 109) with an angle greater than 60 degrees relative to the heart surface (Fig. 5C) or parallel orientation (25%, n = 109) with a spindle angle less than 30 degrees (Fig. 5D). However, cardiomyocytes in B1KO undergo a significantly different division pattern and most of the divisions are parallel with 47% divisions being parallel and 19% being perpendicular (p<0.01, n=102) (Fig. 5E&F). The parallel division pattern may partially explain the thicker compact zone and the absence of trabecula.

**Figure 5:**
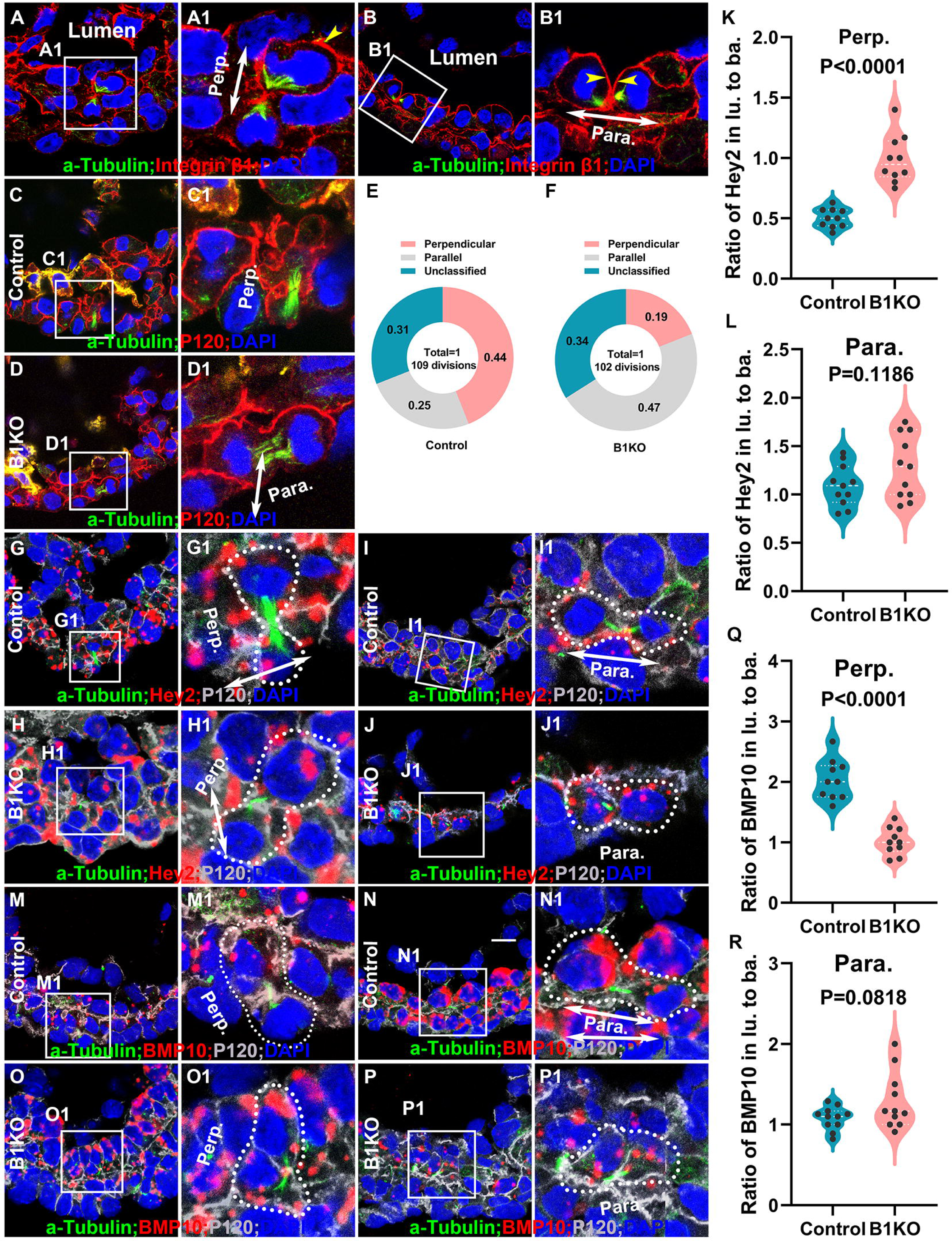
β1 integrins are required for cardiomyocyte-oriented cell division and asymmetric cell division. (A&B) Integrin β1 is asymmetrically distributed in dividing cardiomyocytes, with the luminal side of the daughter cells expressing more Integrin β1 in both perpendicular and parallel divisions. (C-F) Oriented cell division patterns showed the difference between control and B1KO hearts, with a higher percentage of perpendicular divisions in control hearts but a higher percentage of parallel divisions in B1KO hearts. (G-R) In the dividing cardiomyocytes of control hearts, *Hey2* or *BMP10* was asymmetrically distributed to two daughter cells during perpendicular divisions at telophase or at two cell stage (G, M, K, Q), and was symmetrically distributed to two daughter cells during parallel divisions at telophase or at two cell stage (I, N, L, R), but in the B1KO hearts, *Hey2* or *BMP10* was symmetrically distributed to two daughter cells during both perpendicular (H, O, K, Q) and parallel (J, P, L, R) divisions. Scale bar: 10μm.

Perpendicular-oriented cell division is an extrinsic asymmetric cell division that contributes to trabecular specification and is a mechanism that causes the trabecular cardiomyocytes to be distinct from the cardiomyocytes in the compact zone^12^. The cues that regulate asymmetric cell division are unknown. The asymmetric distribution of β1 and trabecular specification defects in B1KO suggest that β1 integrins may regulate asymmetric cell division. We wished to examine the distribution of mRNAs in the dividing cells at telophase, or the two daughter cells when cytokinesis is completed but the midbody is still present^12^. E8.75 heart sections were stained with acetylated α-Tubulin and p120 and hybridized with probes to *Bmp10* or *Hey2* mRNA using RNAScope. We counted the number of mRNA signal dots in dividing cells and quantified the ratio between the two domains of the dividing cell at telophase or the two daughter cells after the division. We found that the *Hey2* mRNAs display asymmetric distribution between daughter cells resulting from perpendicular divisions as the ratios of *Hey2* in luminal daughter to abluminal daughter are close to 0.5 (Fig. 5G, G1 &K), but not between the daughter cells resulting from parallel division (Fig. 5I, I1 &L). This suggests that the different geometric locations or the different signals that the two daughter cells receive cause asymmetric levels of mRNAs. This phenomenon is known as an extrinsic asymmetric cell division and has been well-documented previously in *Drosophila* ovarian stem cells ^44^. In B1KO hearts, *Hey2* did not display an asymmetric distribution between the two daughter cells resulting from perpendicular or parallel cell division (Fig. 5H, 1, J, J1, K &L). We found that *Bmp10* was asymmetrically distributed between the two daughter cells resulting from asymmetric cell division, with the ratios of *Bmp10* in luminal daughter to abluminal daughter being close to 2 (Fig. 5 M, M1 &Q). In parallel divisions, the asymmetric distribution of *Bmp10* is not observed (Fig. 5 N, N1 &R). In B1KO hearts, *Bmp10* did not display an asymmetric distribution between the two daughter cells resulting from either perpendicular or parallel cell division (Fig. 5O, O1, P, P1, Q &R). Therefore, sister cells in all OCD display an equal amount of *Hey2* or *Bmp10* in B1KO, suggesting a failed asymmetric cell division (Fig. 5).

### B1KO hearts display a reduced Notch1 activation, but Notch1 reduction was not sufficient to cause cardiomyocyte organization defect

Multiple signaling pathways, including Notch1 and Nrg1/ErbB, are required for trabecular morphogenesis^45^. To examine if the Notch1 activation was affected in B1KO, Notch1 intracellular domain (N1ICD), a marker for Notch1 activation, was stained in sections from control and B1KO hearts. The percentage of N1ICD+ endocardial cells in B1KO hearts was significantly reduced compared to that of littermate control hearts (Fig. 6A-C). The reduced Notch1 activation was confirmed by Western blot using the whole ventricles of control and B1KO hearts at E9.5 (Fig. 6D&E). pErbB2, a readout for Nrg/ErbB signaling, was not reduced based on pErbB2 western blot using the whole ventricle at E9.5 (Fig. 6F&G).

**Figure 6:**
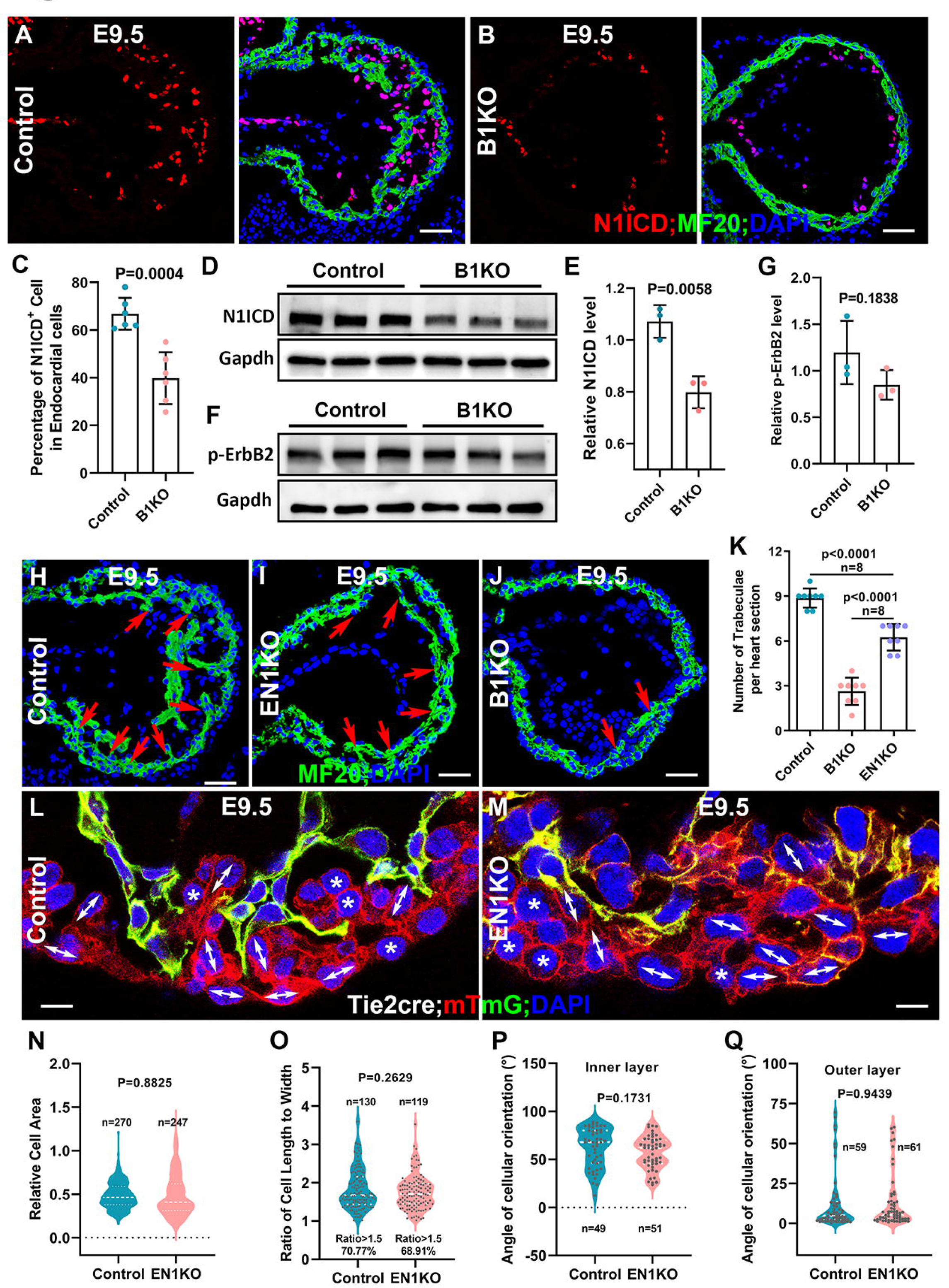
B1KO hearts display reduced Notch1 activation, but Notch1 reduction did not cause a cardiomyocyte organization defect. (A-E) Notch1 activation was reduced in the B1KO hearts, based on N1ICD IF staining (A-C) and Western blot (D&E). (F&G) Nrg/ErbB signaling was not changed in the B1KO hearts, indicated by the unchanged protein level of its readout, p-ErbB2. (H-K) Endothelial-specific Notch1 knockout (EN1KO) hearts mediated by *Tie2-cre* displayed trabeculae growth defect, but the trabecular number per section of EN1KO heart is significantly greater than the B1KO hearts, indicating that the reduction of Notch1 activation is not the reason of the trabeculae initiation defect in B1KO hearts. (L-Q) The cellular shape of cardiomyocytes was examined and quantified between the control and EN1KO heart, their cellular areas (N), percentage of oriented cells (O), and cellular orientation of both the inner and outer layer of the myocardium (P, Q) was not significantly different. Scale bar: 50μm in A&M, H-J, 5μm in L&M.

To directly test whether Notch1 reduction is sufficient to cause trabecular initiation defects and cellular organization defects observed in B1KO hearts, Notch1 was deleted via *Tie2-Cre* to generate endothelial specific Notch1 knockout (EN1KO), and the trabecular initiation and cellular organization were examined. We found that the EN1KO hearts displayed trabeculation defects and died before E10.5, but the number of trabeculae per section in EN1KO is significantly more than the B1KO (Fig. 6H-K). The cellular areas and cellular orientation patterns in inner and outer layer of the myocardium were compared between control and EN1KO and found to be not different (Fig. 6 L-Q). This suggests that cellular orientation and organization are not affected in the EN1KO hearts and the reduced Notch1 activation is not likely to be responsible for the cellular organization and distribution defects in B1KO.

### β1 integrins regulate cellular behaviors and organization autonomously

We applied a mosaic model to determine if the effects of *Itgb1* deletion on cellular organization and behaviors are cell-autonomous or non-autonomous. We employed the inducible sparse lineage tracing system and the embryo clearing system to genetically label, trace, image, and analyze individual clones at single cell levels ^46–48^. The improved imaging depth and scale of the cleared hearts allow for comprehensive 3D reconstruction of the heart and analysis of a single clone with spatial detail at the whole-heart scale. This approach enables us to infer the division patterns and cellular behaviors during trabecular morphogenesis^12,49^. Specifically, we crossed *ROSA26-Cre^ERT^*^2^ (iCre); *Itgb1^fl/+^*, in which iCre nuclear localization can be induced by tamoxifen^50^, with the reporter female mouse *ROSA26-Confetti* (*Conf*); *Itgb1^fl/fl^* or *mTmG*; *Itgb1^fl/fl^*. mTmG reporter was only used to determine the cell shape and cell size. *Conf* reporter mice can stochastically generate nuclear green, cytoplasmic yellow, cytoplasmic red, or membrane-bound blue cells upon Cre mediated recombination^12,46^. Tamoxifen at a concentration of 20-μg/gram-body weight was delivered to pregnant females via gavage when embryos were at E7.75, a stage when the myocardium is a monolayer, and the trabeculae are not yet initiated. 72 hours after induction, single labeled cells have undergone several rounds of cell division to exhibit specific geometric patterns^12^. Based on the geometric distribution and anatomical annotation of each clone, the clones were categorized into three different patterns as previously reported^12^: 1.) Surface or Parallel clones, in which most of the cells localize to the outer layer of myocardium and likely are derived from parallel division; 2.) Transmural or Perpendicular clones, in which the cells localize to both compact and trabecular zones with only one or two cells remaining in the outer compact zone; 3) Other unclassified clones, in which all the cells of the clone localize to the trabecular zone and likely are derived from directional migration of the labeled cell. We compared the clonal patterns between control (*iCre;Itgb1^fl/+^; mTmG)* and iKO, and found that 44% of the control clones are perpendicular (Fig. 7A), and 51% of the iKO clones are parallel (Fig. 7B). The clonal patterns between control and iKO clones were significantly different based on a chi-square test (Fig. 7C&D). iKO clones display a higher percentage of surface clones and lower percentages of transmural and trabecular clones, suggesting that the β1 integrins have roles in OCD and directional migration. We also examined the migration depth via the whole heart clearing and 3-dimensional (3D) imaging^12^. The migration depth is defined by the distance from the innermost cell of the clone to the myocardial surface. We found that the iKO clones migrated a shorter distance compared to the control clones (Fig. 7E). We also measured the number of cells in each clone and found that the number of cells of the control clones is significantly larger than the iKO clones (Fig. 7F). These data suggest that β1 integrins regulate cardiomyocyte migration and proliferation in an autonomous manner during trabecular morphogenesis.

**Figure 7:**
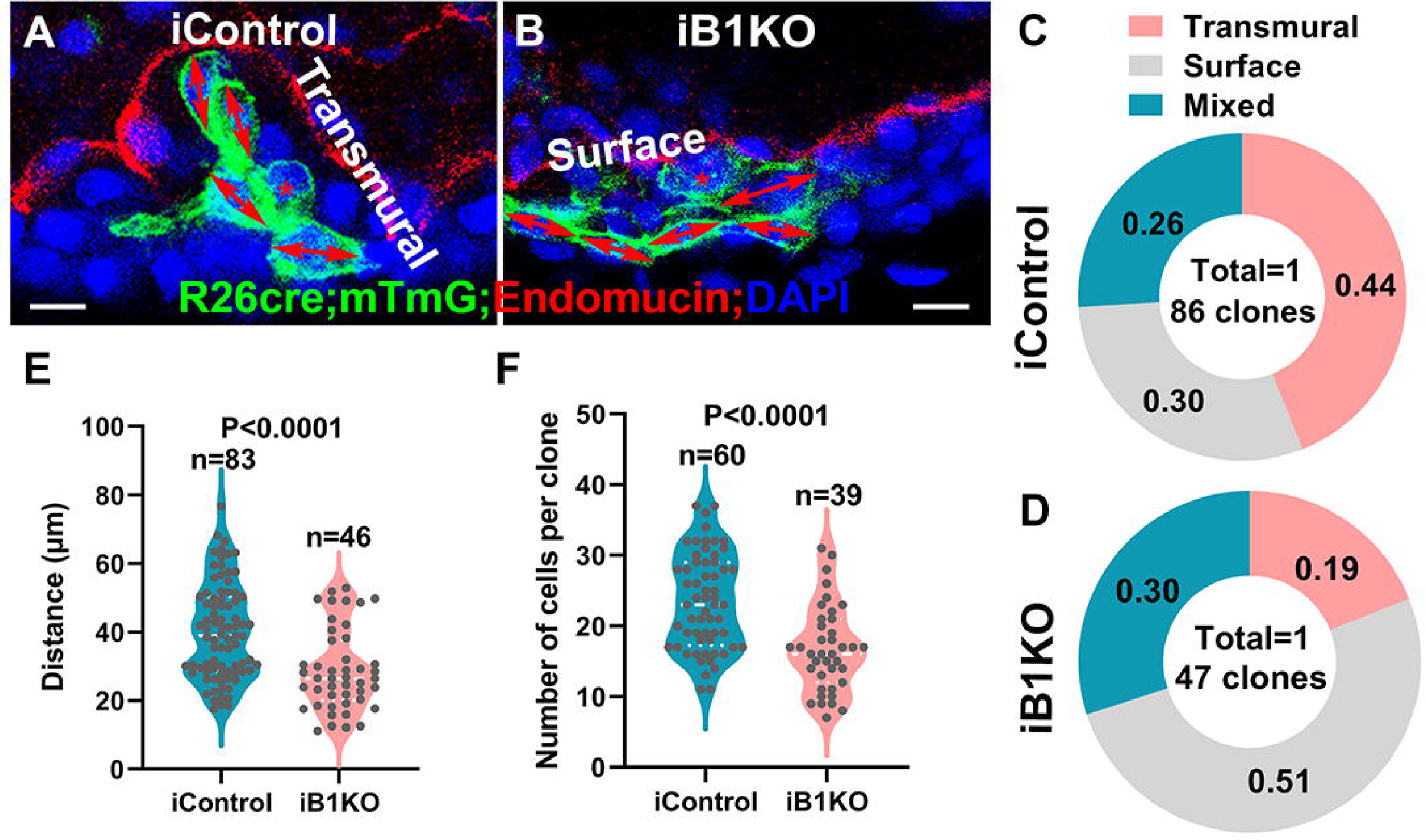
Single-cell lineage tracing shows abnormal cellular behaviors. (A-D) iControl hearts showed a significantly higher percentage of transmural clones but lower percentage of surface clones than iB1KO hearts. (E) iB1KO clones showed significantly shorter migration distances compared with iControl clones. (F) iB1KO clones displayed a smaller clone size compared with iControl, with fewer cells in one clone. Scale bar: 5μm.

## Discussion

Our study demonstrates that *Itgb1* deletion at an early stage of ventricular wall formation results in distinct cardiac phenotypes that are different from those reported in previous studies in which *Itgb1* was deleted at later stages^25,51–56^. Our study shows that β1 integrins are required at early stage for cardiomyocytes to attach to the ECM network, and that this engagement provides structural support for cardiomyocytes to maintain cell shape and undergo cellular behavior that establishes proper cellular organization. Deletion of *Itgb1* leads to disengagement of cardiomyocytes from the ECM network, causing abnormal cellular organization as evidenced by failed trabeculation.

### β1 integr**ins regulate heart formation and function in a spatiotemporal manner**

Integrins are transmembrane heterodimeric proteins and reside at the interface of cardiomyocytes and ECM. These molecules are critical to mediate the cell-cell and ECM interactions that allow for contraction^27^. *Itgb1*, *Itga6,* and *Itga5* are the three most highly expressed integrin subunit genes in the E9.5 heart based on mRNA deep sequencing (Table 1). *Itgb1* is expressed in all types of cells in the heart, and global deletion of *Itgb1* is lethal at the pre-implantation stage^20,21^. Endothelial-specific knockout of *Itgb1* via *Tie2-Cre* results in severe vascular defects and lethality at E10.5^23,24^, and deletion of *Itgb1* via the inducible *VE-Cad^CreERT^*^2^ disrupts endothelial cell polarity and arteriolar lumen formation^22^. Cardiomyocyte-specific knockout of the *Itgb1* via *Mlc-2vCre* results in myocardial fibrosis and cardiac failure in the adult heart^25^, and *Itgb1* deletion via the transgenic *Nkx2.5Cre* line perturbs the trabecular compaction and cardiomyocyte proliferation with progressive cardiac abnormalities seen toward birth^26^. In our study, *Itgb1* deletion via the knockin *Nkx2.5^Cre/+^*, which is expressed in the heart earlier than the transgenic *Nkx2.5Cre* (possibly due to the promoter region of the transgenic *Nkx2.5Cre* not including all the regulatory elements), causes defects in trabecular formation. *Itga6* is enriched in the early heart as evidenced by *in situ* hybridization (ISH) and immunostaining^57^, and the specific expression of α6 in the trabecular zone during compaction^58^ suggests its potential functions in compaction. *Itga6* global knockout dies around birth due to epidermolysis bullosa, and cardiovascular abnormalities were not extensively examined^59^. The *Itga5* global knockouts display a thicker compact zone, as do *Fn1* global knockouts^60^, suggesting a cellular organization defect. Mice homozygous for a null mutation of the *Itgb5* or *Itga1,* or *Itgb3* develop, grow, and reproduce normally^61–64^. The low expression of *Itgav* in the heart is maintained throughout cardiac development^65^. *Itgav* and *Itgb3* integrins were not able to functionally compensate for the loss of *Itgb1* function during specialization and terminal differentiation of cardiomyocytes^30^. *Itga4* deletion cause a defect in epicardial and coronary development^66^. *Itgb4* was not expressed in the heart^57^.

The B1KO hearts display a variety of defects in trabecular formation, specification, compaction, cellular orientation, cellular organization, and proliferation, suggesting that β1 integrins play multiple functions during trabecular morphogenesis. Our results indicate that β1 integrins are essential for ECM organization and extracellular space formation at early stages and the cardiomyocytes attach to the ECM. Our data indicates that β1-mediated cell-ECM interaction at early stage is required to establish cell shape, cellular organization, and tissue architecture, while at later stage these functions may not be essential, as the cell shape, cellular organization, and tissue architecture were already established.

### β1 integrins regulate cellular behaviors and organization during ventricular wall formation

It was exciting to find that β1 was asymmetrically distributed in the myocardium and in the individual cardiomyocytes, with the β1 subunit being enriched in the luminal side of the myocardium and cardiomyocytes at E9.5. Fn, a ligand for α5β1 integrins, is asymmetrically distributed in the myocardium and cardiomyocytes, a similar pattern to that of β1 (Fig. 8A), but also surrounds cardiomyocytes, potentially creating an Fn network for cardiomyocytes. Laminin 411 and collagen IV, two ligands of β1 integrins, are asymmetrically distributed to the luminal side of the myocardium and cardiomyocytes. Furthermore, the ablation of β1 disrupts the localization and protein stability of plasma Fn (Fig. 1F&8B), suggesting that multiple but distinct integrin-ECM interactions are essential for the early cardiac morphogenesis. The asymmetric distributions of β1, Fn, laminin 411, and Collagen IV to the luminal side of the myocardium suggest a polarized myocardium, and this polarized ECM might establish a polarized network frame. The cardiomyocytes attach to the ECM scaffold in the myocardium. The ECMs are enriched with growth factors, provide growth cues for the cells inside, and provide the frame for the cardiomyocytes to be stabilized in the myocardium. β1 integrins from cardiomyocytes engage the cardiomyocytes to the ECM-dependent network frame via its interaction between cardiomyocytes and ECMs (Fig. 8C). This ECM-dependent scaffold provides support for the cellular organization, cellular behaviors, and an ECM niche for cardiomyocyte maturation and ventricular wall specification. The disruption of the interaction between ECMs and cardiomyocytes by deleting *Itgb1* would dis-engage the cardiomyocytes from the ECM scaffold frame, such that the cardiomyocytes would lose their position in the myocardium, their normal cellular behaviors became awry, their cellular organization was disrupted, and fail to initiate and then establish the tissue architecture to form trabeculae. The random cellular organization and lacking growth direction of cells might cause abnormal distribution of cardiomyocytes between the trabecular and compact zone, resulting in reduced trabeculae and a thicker compact zone. These data suggest that β1 integrins are required for cardiomyocyte organization and cellular distribution between trabecular and compact zones. The loss of β1 and separation of cardiomyocyte from the ECM might disrupt the niche that fosters the extrinsic cell division and ventricular wall specification. However, the disruption of cell-ECM interaction in B1KO does not mean the cardiomyocytes can float in the myocardium freely, as N-Cadherin protein level and localization in B1KO was not changed obviously, cell-cell adherens junctions were not affected, and cardiomyocytes are linked to each other in the myocardium based on immunostaining and EM pictures. Phenotypes of Fn knockout mice support this model, as global Fn knockout hearts display a myocardial organization defect with an absence of lumen, and Fn is required for heart morphogenesis^31^.

**Figure 8:**
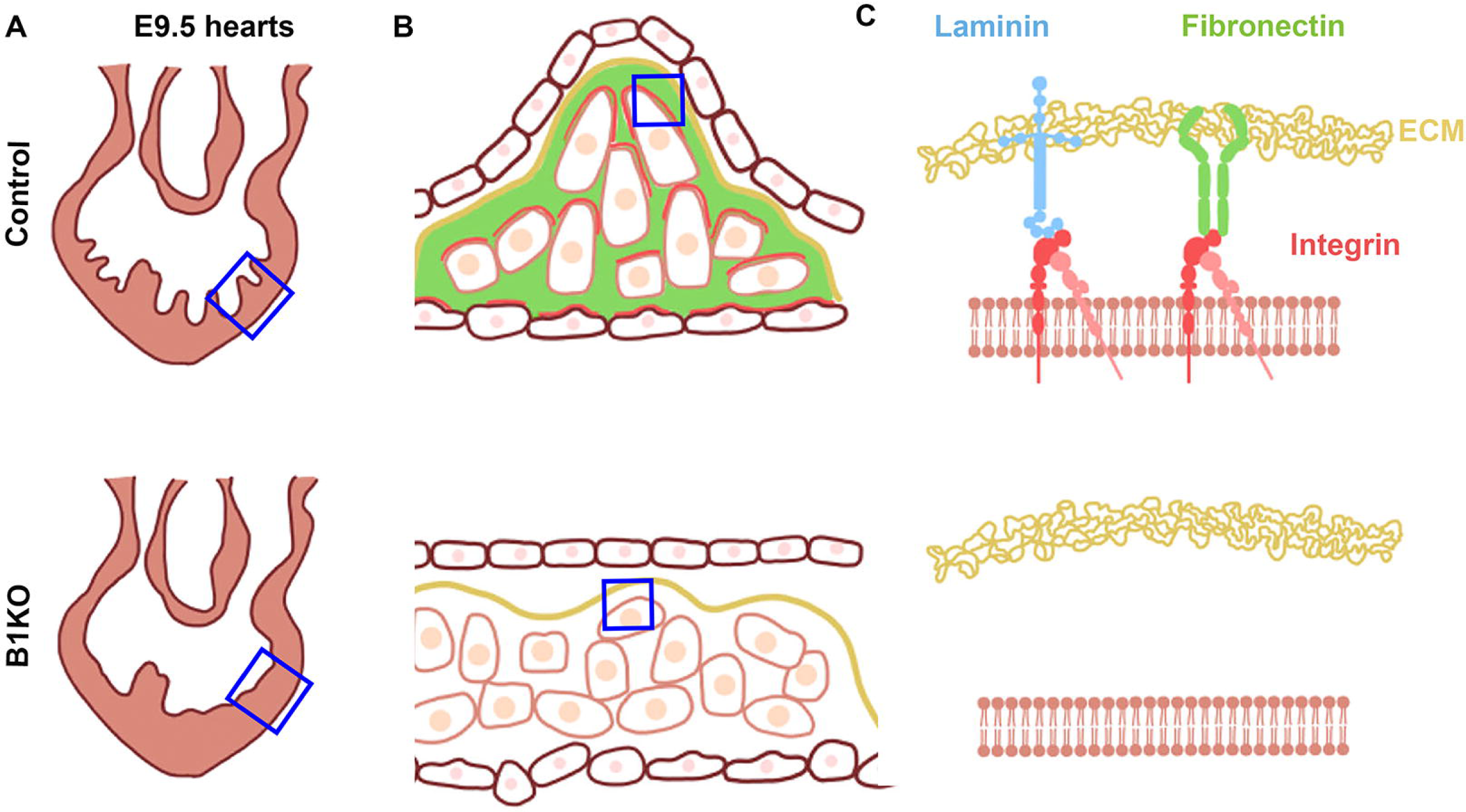
Schematic pictures illustrate the β1 integrins regulating cellular behaviors and organization during ventricular wall formation. (A&B) β1 was asymmetrically distributed in the cardiomyocytes, with being enriched in the luminal side of the cardiomyocytes in E9.5 control heart. (C) The β1, Fn and laminin 411 asymmetrically distributed to the luminal side of the myocardium and establish a polarized network frame, providing a scaffold for the cardiomyocytes to be stabilized in the myocardium. Loss of β1 in the myocardium caused the failure of forming the polarized network frame.

In summary, the ECM-dependent network provides geometrical confinement and a supporting frame for the cardiomyocytes inside the myocardium. The geometrical confinement and supporting frame might shape cardiomyocytes’ proper cellular behavior and organization. β1 integrins play a major role in establishing cell-ECM interaction, which is required to adjust the cell shape in response to signaling and physiological cues to achieve the proper cellular behaviors (Fig. 8). This cell-ECM interaction is required to establish the cardiomyocyte organization or arrangement inside the myocardium. The perturbation of integrin disrupts the correct geometrical confinement and supporting structure, resulting in trabeculation defects.

---

**Suppl. Figure 1. Asymmetrical distribution of Integrin β1 in E8.75 heart.**

(A) Integrin β1 is asymmetrically distributed in dividing cardiomyocytes, with higher enrichment to the membrane to the luminal side. A 10μm-thick section was imaged via a z-stack manner with 0.5μm interval z size (z1-z9). Scale bar: 10μm.

**Suppl. Figure 2. Expression of β1 integrin ligands in the E9.5 hearts.**

(A&B) Immunostaining for Integrin α5 showed that it is expressed in both trabecular and compact cardiomyocytes, and its expression pattern didn’t change in the control hearts compared with B1KO hearts. (C&D) ITGA6 RNAscope revealed that its mRNA level didn’t show a significant difference between control and B1KO hearts based on RNAscope. (E&F) The cellular isoform of Fibronectin is mainly expressed in the AV canal but not in the ventricles by immunostaining, and its expression didn’t change in the B1KO hearts. (G-J) Immunostaining for Laminin 411 and Collagen IV showed that they expressed in the basement membrane and asymmetrically enriched to the luminal side of the myocardium, and their expression didn’t change in the B1KO hearts. Scale bar: 50μm.

**Suppl. Figure 3. B1KO heart displays cardiac progenitor cell differentiation and cardiomyocytes early maturation defects.**

(A&B) The cardiac progenitor cells close to the endoderm express MF20 in the B1KO heart but not in the control heart, indicating the B1KO displayed cardiac progenitor cell differentiation defect. (C) BMP10 mRNA level increased, but the *Hey2* level decreased in the B1KO hearts. (D) The percentage of P21 expressing cardiomyocytes significantly increased in B1KO hearts. (E&F) Sarcomeric array formation in the compact zone of B1KO hearts is more organized than that of the control hearts but less organized than cells in the trabecular zone of the control.

## Acknowledgments

We thank Wu’s lab members for their scientific discussion.

## Author contributions

Conceptualization: MW

Methodology: LM, YL, YX, ARB, AC, CMD, MW

Investigation: LM, YL, YX, MW

Visualization: LM, YL, YX, MW

Funding acquisition: MW, LM, ARB

Project administration: MW

Supervision: MW

Writing – original draft: MW, LM

Writing – review & editing: LM, YL, ARB, AC, CMD, MW

## Data Availability Statement

All data are available in the main text or the supplementary materials. Unpublished data is available upon request. The mouse lines that we generated are available upon request.

## References

1. Olson, E.N. (2004). A decade of discoveries in cardiac biology. Nat Med 10, 467–474. 10.1038/nm0504-467nm0504-467 [pii].

2. Manasek, F.J. (1968). Embryonic development of the heart. I. A light and electron microscopic study of myocardial development in the early chick embryo. Journal of morphology 125, 329–365. 10.1002/jmor.1051250306.

3. Van Mierop, L.H. (1979). Embryology of the univentricular heart. Herz 4, 78–85.

4. Sedmera, D., Pexieder, T., Vuillemin, M., Thompson, R.P., and Anderson, R.H. (2000). Developmental patterning of the myocardium. Anat Rec 258, 319–337.

5. Icardo, J.M., and Fernandez-Teran, A. (1987). Morphologic study of ventricular trabeculation in the embryonic chick heart. Acta Anat (Basel) 130, 264–274.

6. Jenni, R., Rojas, J., and Oechslin, E. (1999). Isolated noncompaction of the myocardium. N Engl J Med 340, 966–967. 10.1056/NEJM199903253401215.

7. Breckenridge, R.A., Anderson, R.H., and Elliott, P.M. (2007). Isolated left ventricular non-compaction: the case for abnormal myocardial development. Cardiol Young 17, 124–129. 10.1017/S1047951107000273.

8. Weiford, B.C., Subbarao, V.D., and Mulhern, K.M. (2004). Noncompaction of the ventricular myocardium. Circulation 109, 2965–2971. 10.1161/01.CIR.0000132478.60674.D0.

9. Zhang, W., Chen, H., Qu, X., Chang, C.P., and Shou, W. (2013). Molecular mechanism of ventricular trabeculation/compaction and the pathogenesis of the left ventricular noncompaction cardiomyopathy (LVNC). Am J Med Genet C Semin Med Genet 163, 144–156. 10.1002/ajmg.c.31369.

10. Samsa, L.A., Yang, B., and Liu, J. (2013). Embryonic cardiac chamber maturation: Trabeculation, conduction, and cardiomyocyte proliferation. Am J Med Genet C Semin Med Genet 163, 157–168. 10.1002/ajmg.c.31366.

11. Wu, M. (2018). Mechanisms of Trabecular Formation and Specification During Cardiogenesis. Pediatr Cardiol. 10.1007/s00246-018-1868-x.

12. Li, J., Miao, L., Shieh, D., Spiotto, E., Li, J., Zhou, B., Paul, A., Schwartz, R.J., Firulli, A.B., Singer, H.A., et al. (2016). Single-Cell Lineage Tracing Reveals that Oriented Cell Division Contributes to Trabecular Morphogenesis and Regional Specification. Cell reports 15, 158–170. 10.1016/j.celrep.2016.03.012.

13. Wu, M., and Li, J. (2015). Numb family proteins: novel players in cardiac morphogenesis and cardiac progenitor cell differentiation. Biomolecular concepts 6, 137–148. 10.1515/bmc-2015-0003.

14. Zhao, C., Guo, H., Li, J., Myint, T., Pittman, W., Yang, L., Zhong, W., Schwartz, R.J., Schwarz, J.J., Singer, H.A., et al. (2014). Numb family proteins are essential for cardiac morphogenesis and progenitor differentiation. Development 141, 281–295. 10.1242/dev.093690.

15. Miao, L., Li, J., Li, J., Tian, X., Lu, Y., Hu, S., Shieh, D., Kanai, R., Zhou, B.Y., Zhou, B., et al. (2018). Notch signaling regulates Hey2 expression in a spatiotemporal dependent manner during cardiac morphogenesis and trabecular specification. Sci Rep 8, 2678. 10.1038/s41598-018-20917-w.

16. Jimenez-Amilburu, V., Rasouli, S.J., Staudt, D.W., Nakajima, H., Chiba, A., Mochizuki, N., and Stainier, D.Y.R. (2016). In Vivo Visualization of Cardiomyocyte Apicobasal Polarity Reveals Epithelial to Mesenchymal-like Transition during Cardiac Trabeculation. Cell reports 17, 2687–2699. 10.1016/j.celrep.2016.11.023.

17. Hynes, R.O. (2002). Integrins: bidirectional, allosteric signaling machines. Cell 110, 673–687.

18. Giancotti, F.G., and Ruoslahti, E. (1999). Integrin signaling. Science 285, 1028–1032.

19. Ridley, A.J., Schwartz, M.A., Burridge, K., Firtel, R.A., Ginsberg, M.H., Borisy, G., Parsons, J.T., and Horwitz, A.R. (2003). Cell migration: integrating signals from front to back. Science 302, 1704–1709. 10.1126/science.1092053.

20. Fassler, R., and Meyer, M. (1995). Consequences of lack of beta 1 integrin gene expression in mice. Genes Dev 9, 1896–1908.

21. Stephens, L.E., Sutherland, A.E., Klimanskaya, I.V., Andrieux, A., Meneses, J., Pedersen, R.A., and Damsky, C.H. (1995). Deletion of beta 1 integrins in mice results in inner cell mass failure and peri-implantation lethality. Genes Dev 9, 1883–1895.

22. Zovein, A.C., Luque, A., Turlo, K.A., Hofmann, J.J., Yee, K.M., Becker, M.S., Fassler, R., Mellman, I., Lane, T.F., and Iruela-Arispe, M.L. (2010). Beta1 integrin establishes endothelial cell polarity and arteriolar lumen formation via a Par3-dependent mechanism. Dev Cell 18, 39–51. S1534-5807(09)00488-2 [pii] 10.1016/j.devcel.2009.12.006.

23. Carlson, T.R., Hu, H., Braren, R., Kim, Y.H., and Wang, R.A. (2008). Cell-autonomous requirement for beta1 integrin in endothelial cell adhesion, migration and survival during angiogenesis in mice. Development 135, 2193–2202. 10.1242/dev.016378.

24. Lei, L., Liu, D., Huang, Y., Jovin, I., Shai, S.Y., Kyriakides, T., Ross, R.S., and Giordano, F.J. (2008). Endothelial expression of beta1 integrin is required for embryonic vascular patterning and postnatal vascular remodeling. Mol Cell Biol 28, 794–802. 10.1128/MCB.00443-07.

25. Shai, S.Y., Harpf, A.E., Babbitt, C.J., Jordan, M.C., Fishbein, M.C., Chen, J., Omura, M., Leil, T.A., Becker, K.D., Jiang, M., et al. (2002). Cardiac myocyte-specific excision of the beta1 integrin gene results in myocardial fibrosis and cardiac failure. Circ Res 90, 458–464.

26. Ieda, M., Tsuchihashi, T., Ivey, K.N., Ross, R.S., Hong, T.T., Shaw, R.M., and Srivastava, D. (2009). Cardiac fibroblasts regulate myocardial proliferation through beta1 integrin signaling. Dev Cell 16, 233–244. 10.1016/j.devcel.2008.12.007.

27. Israeli-Rosenberg, S., Manso, A.M., Okada, H., and Ross, R.S. (2014). Integrins and integrin-associated proteins in the cardiac myocyte. Circ Res 114, 572–586. 10.1161/CIRCRESAHA.114.301275.

28. Lockhart, M., Wirrig, E., Phelps, A., and Wessels, A. (2011). Extracellular matrix and heart development. Birth Defects Res A Clin Mol Teratol 91, 535–550. 10.1002/bdra.20810.

29. Gibbs, B.C., Shenje, L., Andersen, P., Miyamoto, M., and Kwon, C. (2018). beta1-integrin is a cell-autonomous factor mediating the Numb pathway for cardiac progenitor maintenance. Biochem Biophys Res Commun 500, 256–260. 10.1016/j.bbrc.2018.04.054.

30. Guan, K., Czyz, J., Furst, D.O., and Wobus, A.M. (2001). Expression and cellular distribution of alpha(v)integrins in beta(1)integrin-deficient embryonic stem cell-derived cardiac cells. J Mol Cell Cardiol 33, 521–532. 10.1006/jmcc.2000.1326.

31. George, E.L., Baldwin, H.S., and Hynes, R.O. (1997). Fibronectins are essential for heart and blood vessel morphogenesis but are dispensable for initial specification of precursor cells. Blood 90, 3073–3081.

32. Sedmera, D., and Thomas, P.S. (1996). Trabeculation in the embryonic heart. Bioessays 18, 607. 10.1002/bies.950180714.

33. Sedmera, D., Reckova, M., DeAlmeida, A., Coppen, S.R., Kubalak, S.W., Gourdie, R.G., and Thompson, R.P. (2003). Spatiotemporal pattern of commitment to slowed proliferation in the embryonic mouse heart indicates progressive differentiation of the cardiac conduction system. Anat Rec A Discov Mol Cell Evol Biol 274, 773–777. 10.1002/ar.a.10085.

34. Kochilas, L.K., Li, J., Jin, F., Buck, C.A., and Epstein, J.A. (1999). p57Kip2 expression is enhanced during mid-cardiac murine development and is restricted to trabecular myocardium. Pediatr Res 45, 635–642.

35. Chen, H., Shi, S., Acosta, L., Li, W., Lu, J., Bao, S., Chen, Z., Yang, Z., Schneider, M.D., Chien, K.R., et al. (2004). BMP10 is essential for maintaining cardiac growth during murine cardiogenesis. Development 131, 2219–2231. 10.1242/dev.01094 dev.01094 [pii].

36. Clay, H., Wilsbacher, L.D., Wilson, S.J., Duong, D.N., McDonald, M., Lam, I., Park, K.E., Chun, J., and Coughlin, S.R. (2016). Sphingosine 1-phosphate receptor-1 in cardiomyocytes is required for normal cardiac development. Dev Biol 418, 157–165. 10.1016/j.ydbio.2016.06.024.

37. Del Monte-Nieto, G., Ramialison, M., Adam, A.A.S., Wu, B., Aharonov, A., D’Uva, G., Bourke, L.M., Pitulescu, M.E., Chen, H., de la Pompa, J.L., et al. (2018). Control of cardiac jelly dynamics by NOTCH1 and NRG1 defines the building plan for trabeculation. Nature 557, 439–445. 10.1038/s41586-018-0110-6.

38. Miao, L., Li, J., Li, J., Lu, Y., Shieh, D., Mazurkiewicz, J.E., Barroso, M., Schwarz, J.J., Xin, H.B., Singer, H.A., et al. (2019). Cardiomyocyte orientation modulated by the Numb family proteins-N-cadherin axis is essential for ventricular wall morphogenesis. Proc Natl Acad Sci U S A 116, 15560–15569. 10.1073/pnas.1904684116.

39. Cherian, A.V., Fukuda, R., Augustine, S.M., Maischein, H.M., and Stainier, D.Y. (2016). N-cadherin relocalization during cardiac trabeculation. Proc Natl Acad Sci U S A 113, 7569–7574. 10.1073/pnas.1606385113.

40. Fassler, R., Rohwedel, J., Maltsev, V., Bloch, W., Lentini, S., Guan, K., Gullberg, D., Hescheler, J., Addicks, K., and Wobus, A.M. (1996). Differentiation and integrity of cardiac muscle cells are impaired in the absence of beta 1 integrin. J Cell Sci 109 (Pt 13), 2989–2999.

41. Liu, J., Bressan, M., Hassel, D., Huisken, J., Staudt, D., Kikuchi, K., Poss, K.D., Mikawa, T., and Stainier, D.Y. (2010). A dual role for ErbB2 signaling in cardiac trabeculation. Development 137, 3867–3875. 137/22/3867 [pii] 10.1242/dev.053736.

42. Passer, D., van de Vrugt, A., Atmanli, A., and Domian, I.J. (2016). Atypical Protein Kinase C-Dependent Polarized Cell Division Is Required for Myocardial Trabeculation. Cell reports 14, 1662–1672. 10.1016/j.celrep.2016.01.030.

43. Wu, M., Smith, C.L., Hall, J.A., Lee, I., Luby-Phelps, K., and Tallquist, M.D. (2010). Epicardial spindle orientation controls cell entry into the myocardium. Dev Cell 19, 114–125. 10.1016/j.devcel.2010.06.011.

44. Knoblich, J.A. (2008). Mechanisms of asymmetric stem cell division. Cell 132, 583–597. S0092-8674(08)00208-0 [pii] 10.1016/j.cell.2008.02.007.

45. Grego-Bessa, J., Luna-Zurita, L., del Monte, G., Bolos, V., Melgar, P., Arandilla, A., Garratt, A.N., Zang, H., Mukouyama, Y.S., Chen, H., et al. (2007). Notch signaling is essential for ventricular chamber development. Dev Cell 12, 415–429. 10.1016/j.devcel.2006.12.011.

46. Livet, J., Weissman, T.A., Kang, H., Draft, R.W., Lu, J., Bennis, R.A., Sanes, J.R., and Lichtman, J.W. (2007). Transgenic strategies for combinatorial expression of fluorescent proteins in the nervous system. Nature 450, 56–62. 10.1038/nature06293.

47. Susaki, E.A., Tainaka, K., Perrin, D., Kishino, F., Tawara, T., Watanabe, T.M., Yokoyama, C., Onoe, H., Eguchi, M., Yamaguchi, S., et al. (2014). Whole-brain imaging with single-cell resolution using chemical cocktails and computational analysis. Cell 157, 726–739. 10.1016/j.cell.2014.03.042.

48. Hama, H., Kurokawa, H., Kawano, H., Ando, R., Shimogori, T., Noda, H., Fukami, K., Sakaue-Sawano, A., and Miyawaki, A. (2011). Scale: a chemical approach for fluorescence imaging and reconstruction of transparent mouse brain. Nat Neurosci 14, 1481–1488. 10.1038/nn.2928.

49. Shaikh Qureshi, W.M., Miao, L., Shieh, D., Li, J., Lu, Y., Hu, S., Barroso, M., Mazurkiewicz, J., and Wu, M. (2016). Imaging Cleared Embryonic and Postnatal Hearts at Single-cell Resolution. Journal of visualized experiments : JoVE. 10.3791/54303.

50. Badea, T.C., Wang, Y., and Nathans, J. (2003). A noninvasive genetic/pharmacologic strategy for visualizing cell morphology and clonal relationships in the mouse. J Neurosci 23, 2314–2322.

51. Chen, C., Li, R., Ross, R.S., and Manso, A.M. (2016). Integrins and integrin-related proteins in cardiac fibrosis. J Mol Cell Cardiol 93, 162–174. 10.1016/j.yjmcc.2015.11.010.

52. Nishimura, M., Kumsta, C., Kaushik, G., Diop, S.B., Ding, Y., Bisharat-Kernizan, J., Catan, H., Cammarato, A., Ross, R.S., Engler, A.J., et al. (2014). A dual role for integrin-linked kinase and beta1-integrin in modulating cardiac aging. Aging Cell 13, 431–440. 10.1111/acel.12193.

53. Liang, D., Wang, X., Mittal, A., Dhiman, S., Hou, S.Y., Degenhardt, K., and Astrof, S. (2014). Mesodermal expression of integrin alpha5beta1 regulates neural crest development and cardiovascular morphogenesis. Dev Biol 395, 232–244. 10.1016/j.ydbio.2014.09.014.

54. Li, R., Wu, Y., Manso, A.M., Gu, Y., Liao, P., Israeli, S., Yajima, T., Nguyen, U., Huang, M.S., Dalton, N.D., et al. (2012). beta1 integrin gene excision in the adult murine cardiac myocyte causes defective mechanical and signaling responses. The American journal of pathology 180, 952–962. 10.1016/j.ajpath.2011.12.007.

55. Sengbusch, J.K., He, W., Pinco, K.A., and Yang, J.T. (2002). Dual functions of [alpha]4[beta]1 integrin in epicardial development: initial migration and long-term attachment. J Cell Biol 157, 873–882. 10.1083/jcb.200203075jcb.200203075 [pii].

56. Ross, R.S., and Borg, T.K. (2001). Integrins and the myocardium. Circ Res 88, 1112–1119.

57. Thorsteinsdottir, S., Roelen, B.A., Freund, E., Gaspar, A.C., Sonnenberg, A., and Mummery, C.L. (1995). Expression patterns of laminin receptor splice variants alpha 6A beta 1 and alpha 6B beta 1 suggest different roles in mouse development. Dev Dyn 204, 240–258. 10.1002/aja.1002040304.

58. Hierck, B.P., Poelmann, R.E., van Iperen, L., Brouwer, A., and Gittenberger-de Groot, A.C. (1996). Differential expression of alpha-6 and other subunits of laminin binding integrins during development of the murine heart. Dev Dyn 206, 100–111. 10.1002/(SICI)1097-0177(199605)206:1<100::AID-AJA9>3.0.CO;2-M.

59. Georges-Labouesse, E., Messaddeq, N., Yehia, G., Cadalbert, L., Dierich, A., and Le Meur, M. (1996). Absence of integrin alpha 6 leads to epidermolysis bullosa and neonatal death in mice. Nat Genet 13, 370–373. 10.1038/ng0796-370.

60. Mittal, A., Pulina, M., Hou, S.Y., and Astrof, S. (2013). Fibronectin and integrin alpha 5 play requisite roles in cardiac morphogenesis. Dev Biol 381, 73–82. 10.1016/j.ydbio.2013.06.010.

61. Huang, X., Griffiths, M., Wu, J., Farese, R.V., Jr., and Sheppard, D. (2000). Normal development, wound healing, and adenovirus susceptibility in beta5-deficient mice. Mol Cell Biol 20, 755–759.

62. Lane, N.E., Yao, W., Nakamura, M.C., Humphrey, M.B., Kimmel, D., Huang, X., Sheppard, D., Ross, F.P., and Teitelbaum, S.L. (2005). Mice lacking the integrin beta5 subunit have accelerated osteoclast maturation and increased activity in the estrogen-deficient state. Journal of bone and mineral research : the official journal of the American Society for Bone and Mineral Research 20, 58–66. 10.1359/JBMR.041017.

63. Gardner, H., Kreidberg, J., Koteliansky, V., and Jaenisch, R. (1996). Deletion of integrin alpha 1 by homologous recombination permits normal murine development but gives rise to a specific deficit in cell adhesion. Dev Biol 175, 301–313. 10.1006/dbio.1996.0116.

64. Suryakumar, G., Kasiganesan, H., Balasubramanian, S., and Kuppuswamy, D. (2010). Lack of beta3 integrin signaling contributes to calpain-mediated myocardial cell loss in pressure-overloaded myocardium. Journal of cardiovascular pharmacology 55, 567–573. 10.1097/FJC.0b013e3181d9f5d4.

65. Hirsch, E., Gullberg, D., Balzac, F., Altruda, F., Silengo, L., and Tarone, G. (1994). Alpha v integrin subunit is predominantly located in nervous tissue and skeletal muscle during mouse development. Dev Dyn 201, 108–120. 10.1002/aja.1002010203.

66. Yang, J.T., Rayburn, H., and Hynes, R.O. (1995). Cell adhesion events mediated by alpha 4 integrins are essential in placental and cardiac development. Development 121, 549–560.

