## Supplementary material for "β1 integrins regulate cellular behaviors and cardiomyocyte organization during ventricular wall formation": Material and methods

Materials and Methods

**Mouse lines**

Mouse lines of *Itgb1^fl/fl^*, *Notch1^fl/fl^, Rosa26-mTmG (mTmG)*, *Rosa26Cre^ERT2^* *(iCre)* and *Tie2-Cre* were purchased from Jackson Lab. Dr. Robert Schwartz provided *Nkx2.5^Cre/+^* mice. All animal experiments were approved by the Institutional Animal Care and Use Committee (IACUC) at the University of Houston and Albany Medical College and performed according to the NIH Guide for the Care and Use of Laboratory Animals.

**Immunofluorescence (IF)**

Immunofluorescence (IF) staining was performed as previously described. Briefly, embryos or heart samples were fixed in 4% PFA for 2 h at room temperature or overnight at 4 ◦C. After fixation, and samples were washed with PBS and embedded in OCT. Then the sample blocks were sectioned at 10 µm per section. After sectioning, slides were immersed in PBS for 10 min to remove OCT. The sections were then permeabilized with PBT (0.5% Tween in PBS) (if needed) and then blocked for 30 min with TNB blocking buffer (Perkin Elmer, FP1020, Waltham, MA, USA) at RT. After blocking, the sections were incubated with primary antibodies diluted in blocking buffer overnight at 4 ◦C. Then, the slides were washed with PBT 3 × 10 min at RT, followed by secondary antibodies incubation at RT for 1 h. After secondary antibody incubation, the sections were counterstained and mounted in mounting medium (Vectashield, H-1000-10) for confocal imaging. The following primary antibodies were used: Endomucin (1:100, Santa Cruz, sc-65495), MF20 (1:100, DSHB), PECAM (1:50; BD Pharmingen, 550274), N1ICD (1:50; Cell Signaling, 4147S), P57 (1:200, Abcam, ab75974), Integrin b1 (1:100, Millipore, MAB1997), Integrin a5 (1:100, Thermo, PA5-79529), Integrin a6 (1:100, Thermo, 14-0495-85), Fibronectin (gift from Dr. Paula J. Mckeown-Longo’s lab), Laminin (gift from Dr. Susan Laflamme lab), Collogen IV (1:500, Sigma Aldrich, AB756P), P21 (1:100, Abcam, ab109199), p-Smad 1/5/8 (1:500, Cell Signaling, 9511S), Irx3 (1:100, Santa Cruz, sc-30157), P120 (1:100, Santa Cruz, sc-1101), Acetylated tubulin (1:400, Sigma Aldrich, T6793), N-cadherin (1:200, BD Pharmingen, 610920), HABP (1:100, Millipore, 385911), Versican (1:200, Thermo, PA1-1748A), Cleaved Caspase-3 (1:400, Cell Signaling, 9661S),

**Whole embryo immunofluorescence staining and clearing**

Whole embryos were stained as previously described. Briefly, whole embryos were fixed for 2-4 hours in 4% paraformaldehyde, permeabilized for 2 hours in PBS-Tween 20, blocked with 3% BSA, and then incubated with the primary antibody for ∼24 hours. After three washes, the embryos were incubated with the secondary antibody for 24 hours, followed by three more washes with PBS. After staining, embryos were cleared using RapiClear® 1.52 (SunJin Lab, RC152001).

**Single mRNA molecule in situ hybridization (ISH)**

Single mRNA molecule in situ hybridization (ISH) and Immuno-fluorescent staining (IFS) were performed according to the protocol of the kit RNAscope 2.5 HD (RED) Assay (Advanced Cell Diagnostics, 322360) and our published protocol, which enables the detection of single mRNA molecules. Briefly, after fixation for 24 h, the embryos were frozen and embedded in the OCT compound. Sections of the frozen embedded samples were processed following the protocol from the kit. The mRNA expression level in each cell was determined based on the number of mRNA molecules or signal intensity using the confocal scanned pictures, and three scanned sections for each cell were quantified.

**Imaging**

The following systems were used: for confocal imaging, Zeiss LSM 880-NLO confocal microscope system with an Airyscan detector with FAST module on a Zeiss Axio observer Z1 inverted microscope equipped with an internal spectral QUASAR detector. STED Nanoscopes Leica TCS SP8 STED. Stereo images of the heart or embryos were harvested by a stereoscope (Leica M205 FA).

**Quantitative evaluation of the orientation of the cardiomyocyte division plane**

Orientation of the cardiomyocyte division plane was determined by imaging heart sections or cleared whole hearts (described below). Sections or whole hearts of *Nkx2.5^Cre/+^; Itgb1^fl/fl^; mTmG* and *Nkx2.5^Cre/+^; Itgb1^fl/+^; mTmG* at E9.25 were stained with acetylated α-tubulin to determine the spindle orientation. Z-stack images were acquired using a STED Nanoscopes Leica TCS SP8 STED equipped with multi-photon excitation at 1-3 μm per section. Spindle orientations of left ventricular cardiomyocytes in anaphase or early telophase where both centrosomes and nuclei were in the same focal plane were quantified. The spindle orientation was determined by the angle between the spindle axis, determined by the two centrosomes, and the basement membrane or heart surface reference line. Division planes positioned at 60-90° to the basement membrane were classified as perpendicular, those oriented at 0-30° were classified as parallel, and those oriented between 30-60° were considered non-classified. The data shown are a combination of separate analyses from two investigators.

**Quantitative evaluation of the orientation of cardiomyocytes**

Orientation of the cardiomyocyte plane was determined by imaging heart sections or cleared whole hearts. Cardiomyocytes of *Nkx2.5^Cre/+^; Itgb1^fl/fl^; mTmG* and *Nkx2.5^Cre/+^; Itgb1^fl/+^; mTmG* heart at E9.25 were labeled with membrane GFP, and whether a cell is oriented can be determined by the length to width of the cell. If the ratio of length to width of a cell is greater than 1.5, then this cell is oriented. The cellular orientation was determined by the angle between the longitudinal axis of an oriented cell to the heart surface reference line. Cell orientations with 60-90° angles are classified as perpendicular, those oriented at 0-30° are classified as parallel, and those oriented between 30-60° are considered non-classified. The data shown are a combination of separate analyses from three hearts. Only the cellular orientations of cardiomyocytes from the same regions of hearts from the same litter were compared.

**Western Blot Analysis**

Western blot was performed as previously described. Briefly, E9.5 hearts were

harvested and lysed in RIPA buffer. Protein concentration was determined using the BCA kit (Thermo Fisher, 23225), and equal amounts were run on SDS-PAGE using 4–20% Mini-PROTEAN® TGX™ Precast Protein Gels (Bio-Rad, 4561093) and transferred onto PVDF membranes (GE Healthcare Life Science, 10600023) following standard protocols. The antibodies used in the study include GAPDH (1:1000, Santa Cruz, sc-25778), N1ICD (1:1000; Cell Signaling, 4147S), N-cadherin (1:1000, BD Pharmingen, 610920), p-ErbB2 (1:1000, abcam, ab47262), HABP (1:1000, Millipore, 385911), Versican (1:1000, Thermo, PA1-1748A).

**Inducible lineage tracing and mosaic analysis**

Heterozygous *Rosa^CreERT2^* males were crossed with homozygous *mTmG* or *Confetti* females to produce reporter embryos. Tamoxifen (T5648, Sigma), dissolved in sunflower seed oil (S5007, Sigma), was gavaged to pregnant females 7.75 days after coitus or at a specified time, with a dose of tamoxifen at 100 or 50 μg per gram body weight. The embryos were harvested at the indicated age and used for IFS or whole-mount staining. The clones were imaged and then analyzed for cell count and distance (z) from the heart surface to the innermost cell of the clone.

**mRNA deep sequencing**

Total RNA was isolated from nine E9.5 hearts from both control and B1KO for each experiment. As an indication of quality, the RNA had an integrity number of 8 or greater by Bioanalyzer (Agilent Technology). Samples for mRNA deep sequencing were prepared according to the manufacturer’s protocol (mRNASeq 8-Sample Prep Kit, Illumina). The samples were sequenced by the Microarray Core Facility at the University of Texas, Southwestern Medical Center at Dallas. A HiSeq 2000 system (Illumina) was used for SE-50 sequencing (single-ended 50 bp reads), with over 30×10^6^ ‘reads’ per sample. Basic data analysis was performed with CLC-Biosystems Genomic Workbench analysis programs to generate quantitative data for all genes. The quality filtered and trimmed reads were aligned to an annotated mouse reference genome downloaded from the Ensembl Genome Browser. cDNA fragments were mapped back to individual transcripts. After normalization, the RNA-Seq fragment count was used to measure the relative abundance of transcripts. CLC BIO measured transcript abundances in reads per kilobase of transcript per million mapped reads (RPKM). The experiment was repeated three times. Relative expression levels of genes (ratio of B1KO to control) with P<0.05 were considered significantly different.

**Electron microscopy imaging**

Embryos at E9.5 were processed for serial block-face scanning electron microscopy (SBF-SEM). After isolating the embryos at the desired age, samples were kept in a fixative solution (0.1M sodium cacodylate buffer containing 2.5% glutaraldehyde and 2Mm calcium chloride) for 2 hours at room temperature while gently agitating, then keep it at 4 degrees overnight. Then the embryos were washed in a washing buffer (0.1 M sodium cacodylate buffer containing 2 mM calcium chloride) three times for 10 min each at room temperature. Then the embryos were stained with heavy metals (Fe, OsO4, uranyl acetate, lead) before dehydration through an acetone series and embedded in Embed 812 resin (Electron Microscopy Sciences, USA) containing Ketjenblack EC600JD (Lion Specialty Chemicals Co., Japan). The resin-embedded blocks were sputter-coated with gold to further reduce charging during block-face imaging. Tissue blocks were serially sectioned at 100 nm steps using a Gatan 3View2 microtome (Gatan, USA) mounted to a Mira 3 scanning electron microscope (Tescan, USA). Back scatter electron (BSE) detection was used to image the block-face. Serial imaging was conducted under high vacuum (0.047 Pa) using a Schottky emitter, and an accelerating voltage of 9.0 keV Beam intensity ranged from 5-7 on a scale ranging from 1-20, with a pixel dwell time of 32 μs, and a spot size of 4-7 nm. Magnification ranged from 500-36,900x and pixel size from 7.32-542 nm. Subsequent three-dimensional segmentation and reconstruction were conducted using Amira 6.0.1 software (FEI Company, Hillsboro, OR).

**Statistics analysis**

Data are shown as mean ± standard deviation. As specified, an unpaired, two-tailed Student’s t-test and chi-square test were used for statistical comparison. A P-value of 0.05 or less was considered statistically significant.
