## Supplementary figures and images for "β1 integrins regulate cellular behaviors and cardiomyocyte organization during ventricular wall formation"

### Suppl. Fig. 1

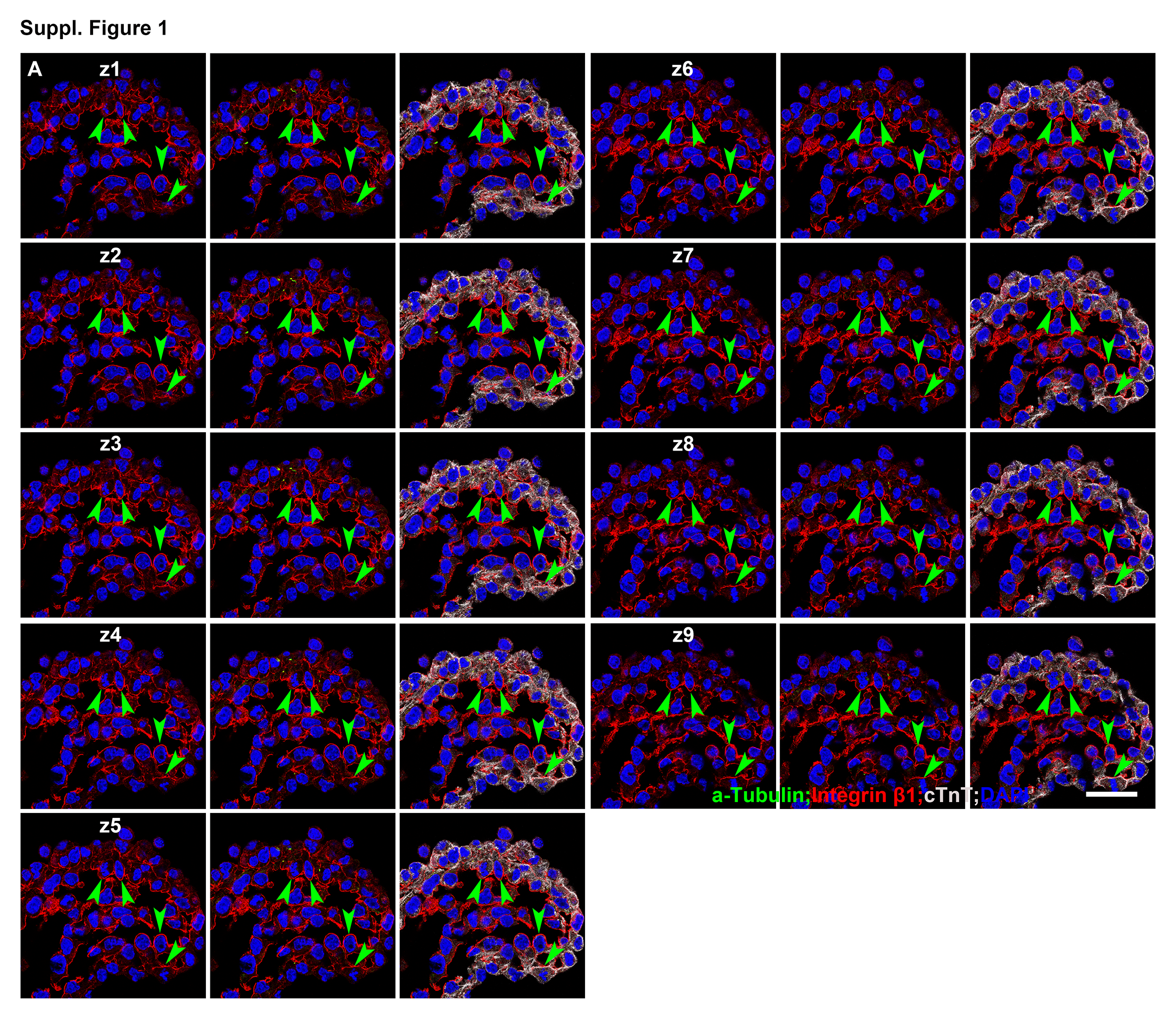

### Suppl. Fig. 2

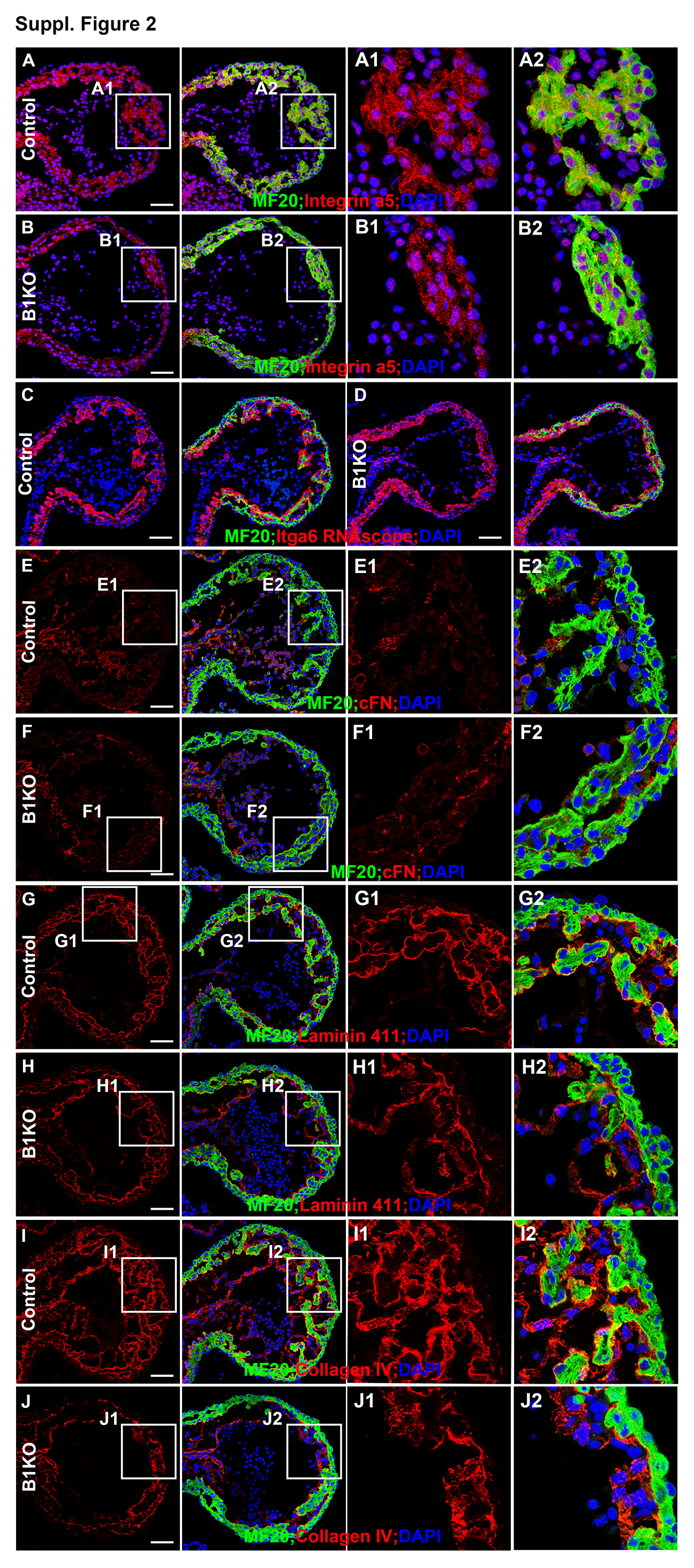

### Suppl. Fig. 3

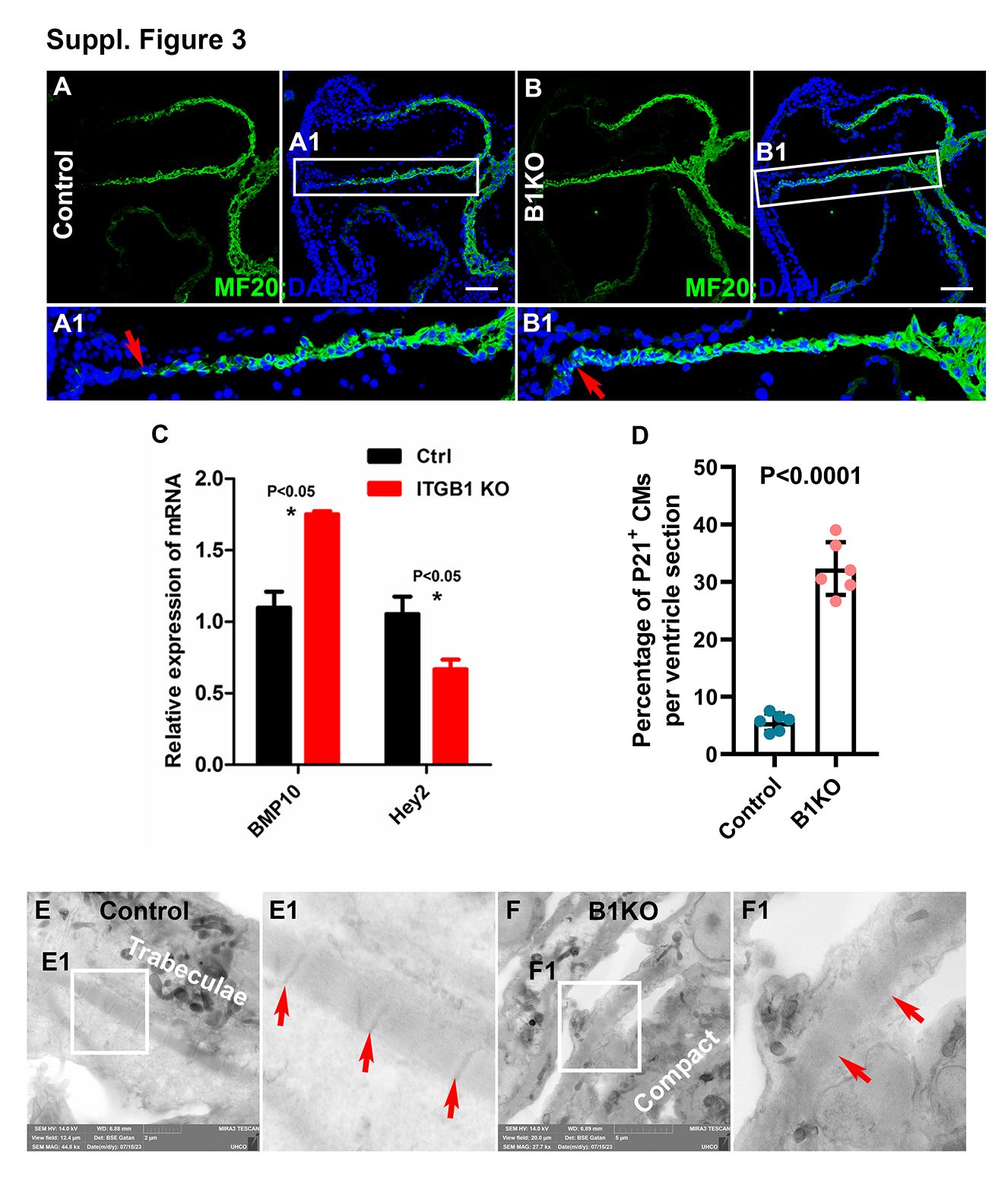
